# ANKLE1 cleaves mitochondrial DNA and contributes to cancer risk by promoting apoptosis resistance and metabolic dysregulation

**DOI:** 10.1101/2021.10.27.466184

**Authors:** Piotr Przanowski, Róża K. Przanowska, Michael J. Guertin

## Abstract

Alleles within the chr19p13.1 locus are associated with increased risk of both ovarian and breast cancer and increased expression of the *ANKLE1* gene. ANKLE1 is molecularly characterized as an endonuclease that efficiently cuts branched DNA and shuttles between the nucleus and cytoplasm. However, the role of ANKLE1 in mammalian development and homeostasis remains unknown. In normal development ANKLE1 expression is limited to the erythroblast lineage and we found that ANKLE1’s role is to cleave the mitochondrial genome during erythropoiesis. We show that ectopic expression of ANKLE1 in breast epithelial-derived cells leads to genome instability and mitochondrial DNA (mtDNA) cleavage. mtDNA degradation then leads to mitophagy and causes a shift from oxidative phosphorylation to glycolysis (Warburg effect). Moreover, mtDNA degradation activates STAT1 and expression of epithelial-mesenchymal transition (EMT) genes. Reduction in mitochondrial content contributes to apoptosis resistance, which may allow precancerous cells to avoid apoptotic checkpoints and proliferate. These findings provide evidence that ANKLE1 is the causal cancer susceptibility gene in the chr19p13.1 locus and describe mechanisms by which higher ANKLE1 expression promotes cancer risk.

## Introduction

Alleles within the cancer susceptibility locus chr19p13.1 were first found to modify the risk of breast cancer in BRCA1 mutation carriers, triple negative breast cancer (TNBC), and ovarian cancer (Antoniou et al. 2010; Bolton et al. 2010). The authors suggested that *BABAM1* was acting as the causal gene to modify breast and ovarian cancer risk based upon its physical interaction with the BRCA1 protein complex and its expression in ovarian cancer. Integration of GWAS data with expression quantitative trait loci (eQTL) analysis implicated either *ABHD8* and/or Ankyrin repeat and LEM-domain containing protein 1 (*ANKLE1*) as candidate causal genes (Lawrenson et al. 2016). The authors concluded that *ABHD8* was the most plausible causal gene in the locus based upon chromatin conformation assays, deletion analysis, expression data, and cell migration experiments (Lawrenson et al. 2016). Later work integrated GWAS and eQTL analysis, along with evolutionary conservation data, ChIP-seq, and chromatin accessibility data to identify the likely causal variant and causal gene, with a focus on variants that disrupt transcription factor binding (Liu et al. 2017). This work also proposed that a variant within a CCCTC-binding factor (CTCF) binding site reduces the affinity for CTCF and causes *ANKLE1* expression to increase (Liu et al. 2017). This work suggested that *ANKLE1* was the most likely the casual gene within the chr19p13.1 breast and ovarian cancer susceptibility locus.

Few studies have explored the molecular, cellular, and physiological functions of ANKLE1. ANKLE1 was first studied in 2012 as an uncharacterized endonuclease that requires its LEM domain and GIY-YIG motifs for DNA cleavage *in vivo* (Brachner et al. 2012). Interestingly, ANKLE1 has high specificity for cleaving branched DNA (Song et al. 2020). Specificity for branched DNA is consistent with an observation in *C*.*elegans* that its homolog resolves chromatin bridges during late mitosis (Hong et al. 2018a) and is involved in the the regulation of meiotic recombination repair and chromosome segregation (Hong et al. 2018b). However, ANKLE1 is dispensable for resolving chromatin bridges, meiotic recombination, and DNA repair in mice (Braun et al. 2016). *ANKLE1* is primarily expressed in hematopoietic tissues of vertebrates, but *ANKLE1*-deficient mice are viable without any detectable phenotype in hematopoiesis (Braun et al. 2016).

Herein we confirm that ANKLE1 expression is normally limited to the erythroblast lineage and we determine that the developmental role of ANKLE1 is to cleave the mitochondrial genome during erythropoiesis. Although this appears to be the molecular and cellular function of ANKLE1, the biological relevance of this function with regards to organismal development remains unclear. We also explore the undesired role that ectopic expression of ANKLE1 plays in conferring breast cancer risk. We find that ectopic expression of ANKLE1 in epithelial breast cells leads to genome instability, the Warburg effect, and resistance to apoptosis.

## Results

### ANKLE1 is the causal gene for breast and ovarian cancer risk in the chr19p13.1 region

Expression quantitative trait loci (eQTL) data have revolutionized how geneticists identify candidate causal genes from genome-wide association study (GWAS) loci. We integrated the most recent meta-analysis of breast cancer GWAS (Zhang et al. 2020) and Genotype-Tissue Expression (GTEx) project data; we found that *ANKLE1* eQTL variants colocalize with the cancer susceptibility GWAS variants (Figure 1A). In contrast, we found minimal colocalization with other genes in the region (Figure S1A). Statistical colocalization analysis (Wallace 2020) demonstrates that *AN-KLE1* expression and genetic risk of breast cancer share a single casual variant in the locus with a probability of 0.75 (Figure 1B). This finding is consistent with recent integrative genomic analyses (Ferreira et al. 2019; Hoffman et al. 2017; Liu et al. 2017). Alleles that are associated with increased risk of breast and ovarian cancer are associated with high expression of *ANKLE1* (Figure 1C and Figure S1B).

**Fig. 1.**
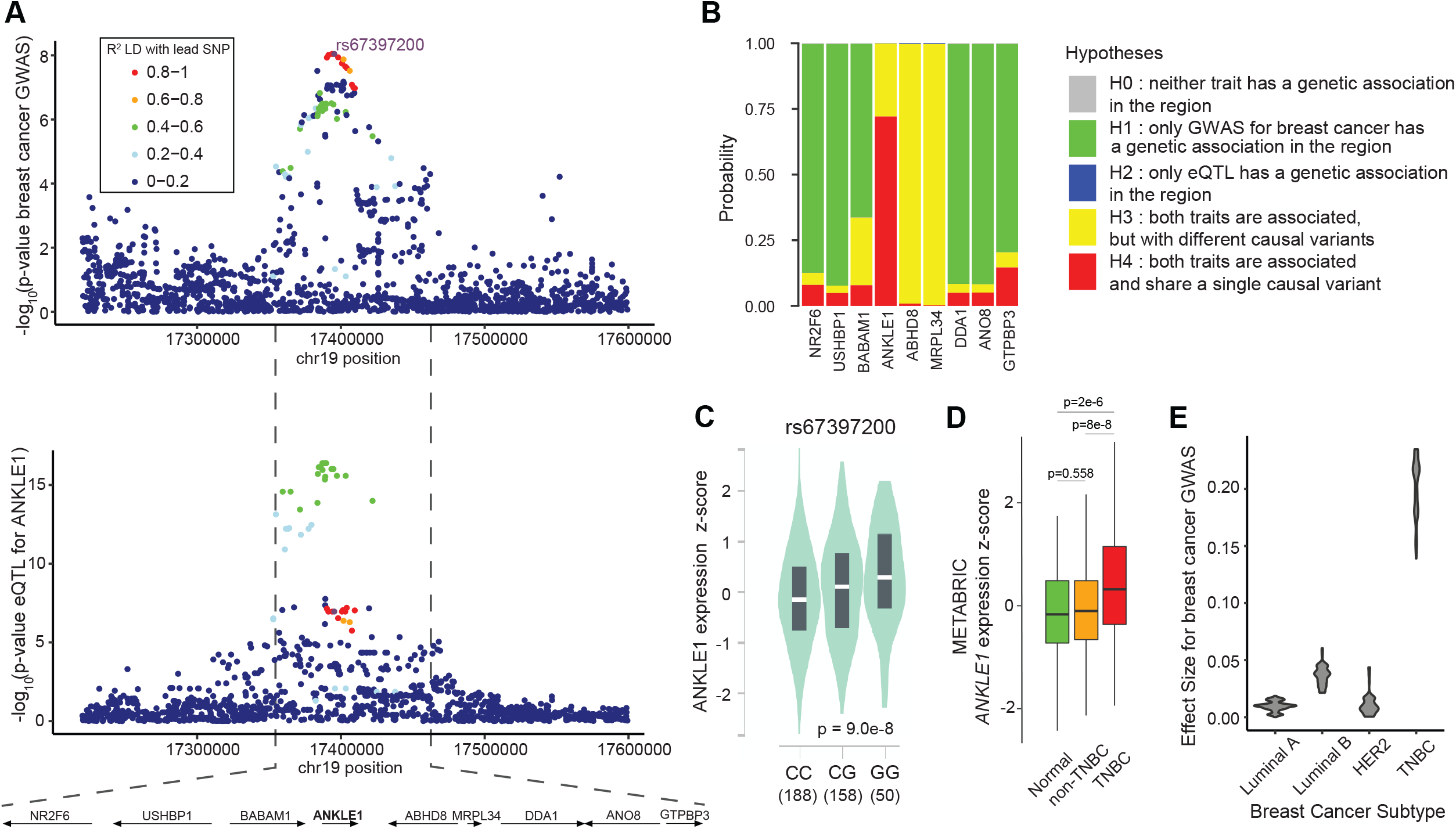
*ANKLE1* is the most likely causal gene for Triple-Negative Breast Cancer risk in the 19p13.1 region. A) The breast cancer susceptibility GWAS variants (upper panel) and *ANKLE1* eQTL variants (lower panel) colocalize. B) Genetic colocalization analysis indicates that both ANKLE1 expression and breast cancer GWAS are associated and likely share a single causal variant. C) The G allele of rs67397200, which is associated with increased breast and ovarian cancer risk, is associated with higher expression of *ANKLE1* in breast tissue. The number of individuals with each genotype is indicated in parentheses. D) *ANKLE1* expression as measured by METABRIC is higher in Triple Negative Breast Cancer (TNBC) than in Normal (tumor-adjacent) or non-TNBC (Luminal A, Luminal B, HER2) tissue (p-values are calculated with a two-tailed t-test). E) Violin plots illustrate that TNBC is the predominant breast cancer subtype associated with the alleles within the 19p13.1 locus.

*ANKLE1* is more highly expressed in triple negative breast cancers (TNBC) compared to adjacent normal tissue and other breast cancer subtypes (Figure 1D). Although we do not claim to identify the causal variant(s) that lead to increased cancer risk, these analyses confirm that ANKLE1 is the most likely causal gene within the region.

Breast cancer is classified into at least four distinct subtypes. The chr19p13.1 locus was originally identified as a breast cancer risk region in BRCA1 mutation carriers and TNBC patients. We compared the effect sizes for the most significant polymorphisms in this region and confirmed that the variants specifically contribute to risk of developing TNBC (Figure 1E). We also compared eQTL significance to disease risk effect size for all genes in the region and these values are most highly correlated with the *ANKLE1* eQTL in TNBC (Figure S1C).

### Expression of ANKLE1 in normal tissue is limited to erythroblasts

Although *ANKLE1* gene expression is associated with breast and ovarian cancer, no Mendelian inherited human diseases are caused by *ANKLE1* mutations and *ANKLE1* knockout mice have no discernible phenotypes (Braun et al. 2016).

In an effort to understand the normal biological role of ANKLE1, we examined ANKLE1 protein expression in different tissues in the Human Protein Atlas (Thul et al. 2017). ANKLE1 protein is only detected in a small subpopulation of cells within the bone marrow tissue (Figure S2A and Figure 2A). Expression analysis of RNA isolated from different bone marrow cell subpopulations indicates that ANKLE1 expression is limited to erythroblasts (Figure 2B) (Konuma et al. 2011). *ANKLE1*’s expression increases during erythroblast differentiation (Figure S2B), which indicates a possible role in red blood cell development. We hypothesized that ANKLE1 promotes enucleation and DNA fragmentation in erythrocytes based on its expression and the presence of its endonuclease domain. We leveraged an *in vitro* model of erythroblast differentiation of human leukemia K562 cells to test this hypothesis (Alves et al. 2011; Witt et al. 2000). Consistent with the expression pattern observed in human erythroblast differentiation (Figure S2B) (Ludwig et al. 2019), *ANKLE1* is transcriptionally activated in differentiating K562 cells (Figure S2C). Contrary to our expectations, DNA fragmentation is indistinguishable throughout differentiation between clonal *ANKLE1* knockout (KO) and clonal wild-type (WT) K562 cells (Figure 2C & Figure S2E&I). This result is consistent with earlier experiments showing that ANKLE1 is dispensable for normal erythropoiesis in *ANKLE1* KO mice (Braun et al. 2016). Lastly, differentiating *ANKLE1* KO clones enucleate at a slightly faster rate (Figure S2F) and contain normal hemoglobin levels (Figure S2G).

**Fig. 2.**
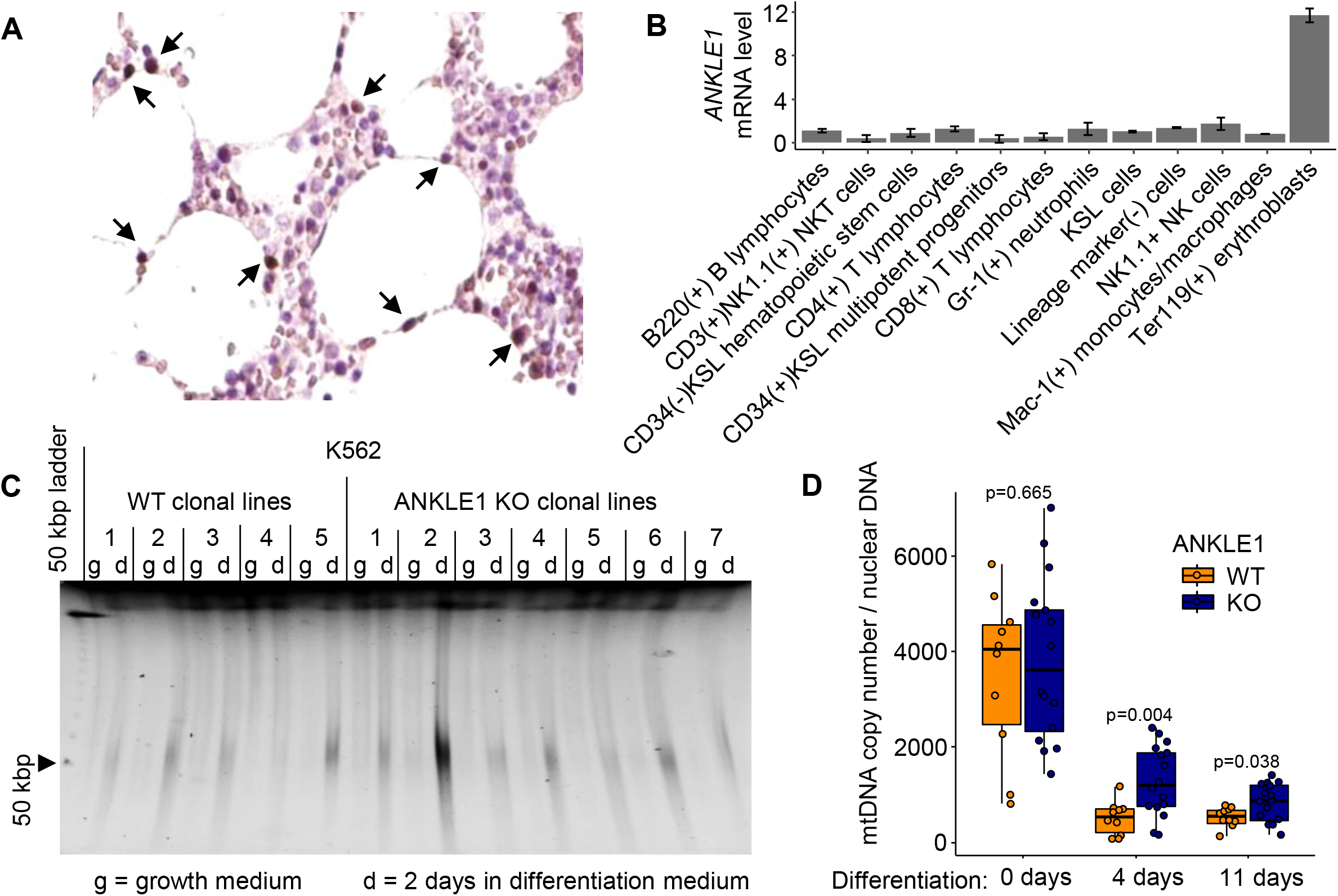
The role of ANKLE1 in development is limited to mtDNA degradation in differentiating erythroblasts. A) Immunohistochemistry (brown) for ANKLE1 in human bone marrow tissue highlights rare population of cells expressing ANKLE1, as shown by the arrows. Images were reproduced with permission from https://www.proteinatlas.org/ENSG00000160117-ANKLE1/tissue/bone+marrow#img. B) Microarray data (GDS3997, probe 1443978at ANKLE1) from human bone marrow shows that *ANKLE1* is specifically expressed in the erythroblast lineage (Konuma et al. 2011). C) Nuclear DNA fragmentation is no different between wild-type (WT) and knock-out (KO) *ANKLE1* K562 cells, as shown by pulse field gel electrophoresis (PFGE) analysis of DNA isolated from undifferentiated and differentiated (2 days) K562 cells. D) Quantitative PCR with primers specific to nuclear and mitochondrial DNA shows that mitochondrial DNA copy number is higher in *ANKLE1* KO K562 clonal lines compared to WT lines during differentiation (p-values are calculated with a two-tailed t-test).

### ANKLE1 localizes to the mitochondria to promote mtDNA degradation and mitophagy

In addition to removing their nucleus, erythrocytes also digest their mitochondria and degrade mtDNA. Mitochondrial DNA is circular, but it is also distinct from nuclear DNA with respect to the lack of nucleosomes and a unique structural feature known as a displacement loop (D-loop) (Kasamatsu et al. 1971). The D-loop structure contains two branched DNA sites; ANKLE1 cleaves branched DNA orders of magnitude more efficiently than B-form DNA (Song et al. 2020). We hypothesized that ANKLE1 functions to cleave mitochondrial DNA in red blood cell development to facilitate mtDNA degradation. We found that mitochondrial DNA copy number is higher in *ANKLE1* KO lines throughout differentiation (Figure 2D). Mitophagy, or autophagy of the mitochondrial organelle, and enucleation occur during erythropoiesis in humans and in the K562 model (Moras et al. 2017) (Figure S2D). Increased mtDNA is not accompanied by an increase in mitochondrial mass (Figure S2H), which suggests that mitophagy is not impaired in *ANKLE1* KO cells.

The *ANKLE1* knockout model of erythropoiesis, mtDNA quantification data, and the preferential cleavage of branched DNA all suggest a role for ANKLE1 in regulating mtDNA abundance through D-loop cleavage. We directly characterized the phenotypes associated with overexpression of ANKLE1 in HEK293T cells. We sorted cells to select for non-apoptotic cells that express either GFP-ANKLE1 or GFP alone (Figure S3A). We found that ANKLE1 expression reduced the levels of mtDNA by over two-fold (Figure 3A) and decreased mitochondrial mass by 25% (Figure 3B). Next, we aimed to determine whether ANKLE1 induces mitophagy by quantifying the colocalization of the autophagasome protein marker LC3 and mitochondria, which is indicative of active mitophagy. We used confocal microscopy to show that mitochondria and autophagasomes stained by either Mito-CFP (an N-terminal mitochondrial localization peptide from COX8) and RFP-LC3 colocalize (Figure 3C). The control GFP expressing cells do not form punctate distributions that colocalize with autophagasomes and mitochondria (Figure 3C and Figure S3B). Moreover, ANKLE1 also colocalizes with staining of lysosome and mitochondria organelles as measures by lysotracker and mitotracker (Figure S3C). The colocalization analyses suggest a mitophagy mediates the decrease in mitochondrial mass, so we blocked autophagasome/lysosome fusion with Bafilomycin A1 to confirm the direct role of autophagy. Bafilomycin A1 inhibits normal turnover of the mitochondria in the control cells and abolishes the effect of ANKLE1-induced decrease in mitochondrial mass (Figure S3D). We further quantified mitophagy by imaging flow cytometry of cells with either GFP or GFP-ANKLE1 that were stained with mitotracker, lysotracker and DAPI (Figure 3D). Bright detail similarity analysis confirmed that ANKLE1 colocalizes with mitochondria (Figure 3E) and, to smaller extent, with lysosomes (Figure 3F). We found that ANKLE1 is uniformly distributed in cytoplasm in some cells, so we compared these cells to cells characterized by punctate ANKLE1 distribution. Cells with punctate distribution of ANKLE1 have reduced mitochondrial mass (Figure S3E&F), which suggests that ANKLE1 translocates to the mitochondria and precipitates mitochondrial degradation. Lastly, mitochondria colocalize with lysosomes more frequently in the presence of ANKLE1, indicating that mitophagy is increased in GFP-ANKLE1 cells compared to GFP-alone (Figure 3G).

**Fig. 3.**
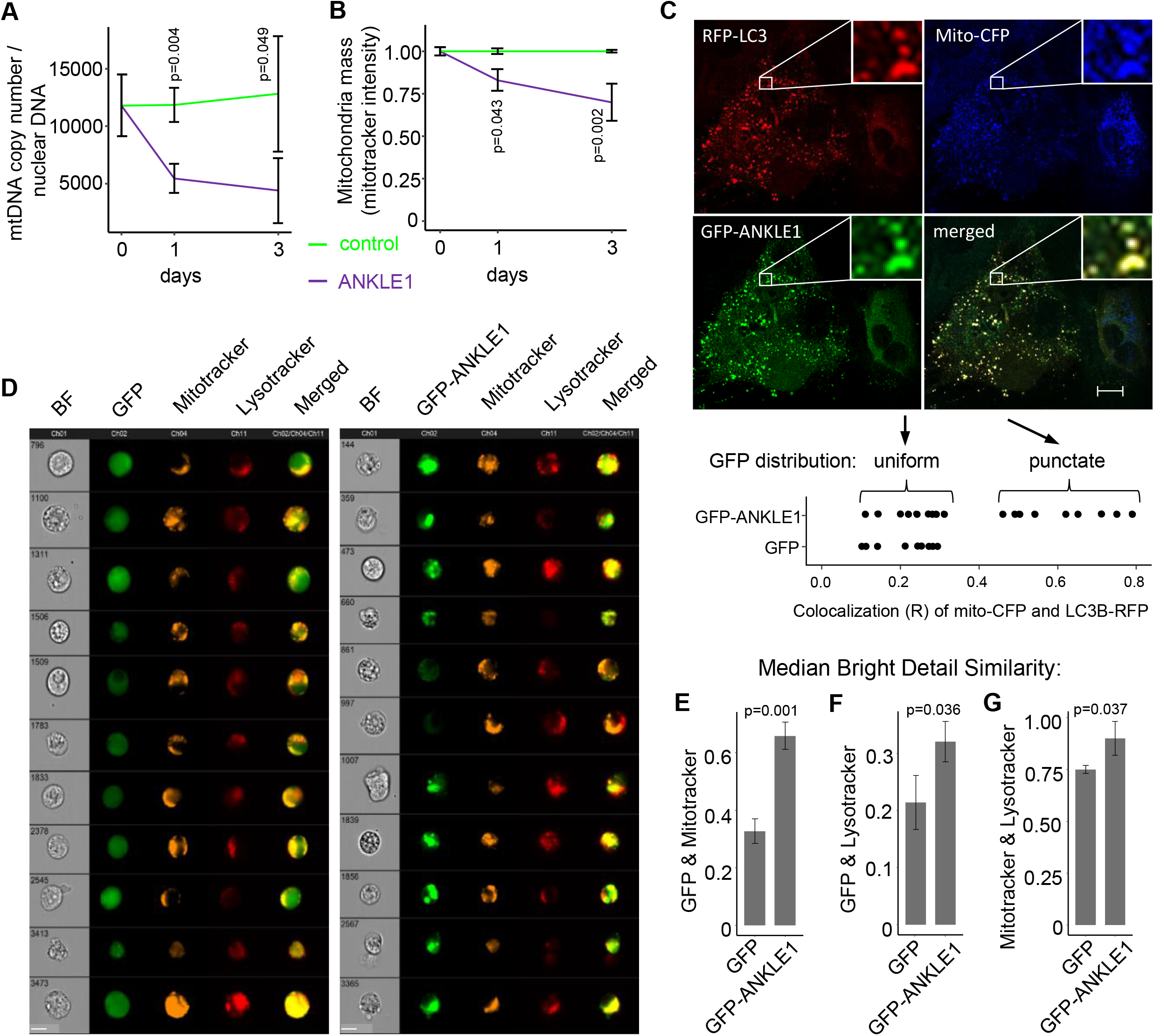
ANKLE1 localizes to mitochondria, degrades mtDNA, and leads to mitophagy. A-B) Overexpression of ANKLE1 decreases mtDNA level (A) and decreases mitochondria mass (B) in HEK293T cells compared to the GFP expressing controls. C) Confocal microscopy imaging (scale bar 10 µm) with fluorescent-tagged mitochondrial targeting sequence present in the N-terminus of COX8, ANKLE1, and LC3 proteins shows colocalization of ANKLE1, mitochondria, and autophagasomes when GFP-ANKLE1 form puncta. The lower portion of the panel quantifies this relationship alongside the GFP control from Figure S3B. D) Representative cells from imaging flow cytometry of HEK293T cells overexpressing GFP (control) or GFP-ANKLE1, stained with DAPI (to exclude dead cells), mitotracker, and lysotracker (scale bar 10 µm) show colocalization of ANKLE1, mitochondria, and lysosomes. E-F) We quantified all the imaging flow cytometry data with bright detail similarity analysis of GFP, mitotracker, and lysotracker which more rigorously shows that ANKLE1 colocalizes with both mitochondria (E) and lysosomes (F). G) The same analysis shows that mitochondria and lysosomes colocalize more in the presence of ANKLE1, indicating an increase in mitophagy (p-values are calculated with a two-tailed t-test).

### Genomic association studies and eQTL data reveal that ANKLE1 expression inversely correlates with mitochondrial DNA copy number

Independent studies found that variants within the chr19p13.1 locus associate with DNA copy number (mtDNA-CN) in human blood (Chong et al. 2022; Ganel et al. 2021; Guyatt et al. 2019), which is consistent with a role of ANKLE1 in regulating mtDNA copy number in erythropoiesis. The same haplotype block that associates with mtDNA-CN is the lead linkage block that associates with breast cancer risk (Figure 4A). Just as ANKLE1 is more highly expressed in TNBC cancers and the effect size for the breast cancer GWAS is amplified in TNBCs (Figure 1D&E), the association between mtDNA-CN and breast cancer risk effect size is unique to the TNBC subtype (Figure 4B). This is consistent with previous observations that TNBC tumors display a higher frequency of mitochondrial defects and lower mtDNA-CN compared to other breast cancer subtypes (Guha et al. 2018).

**Fig. 4.**
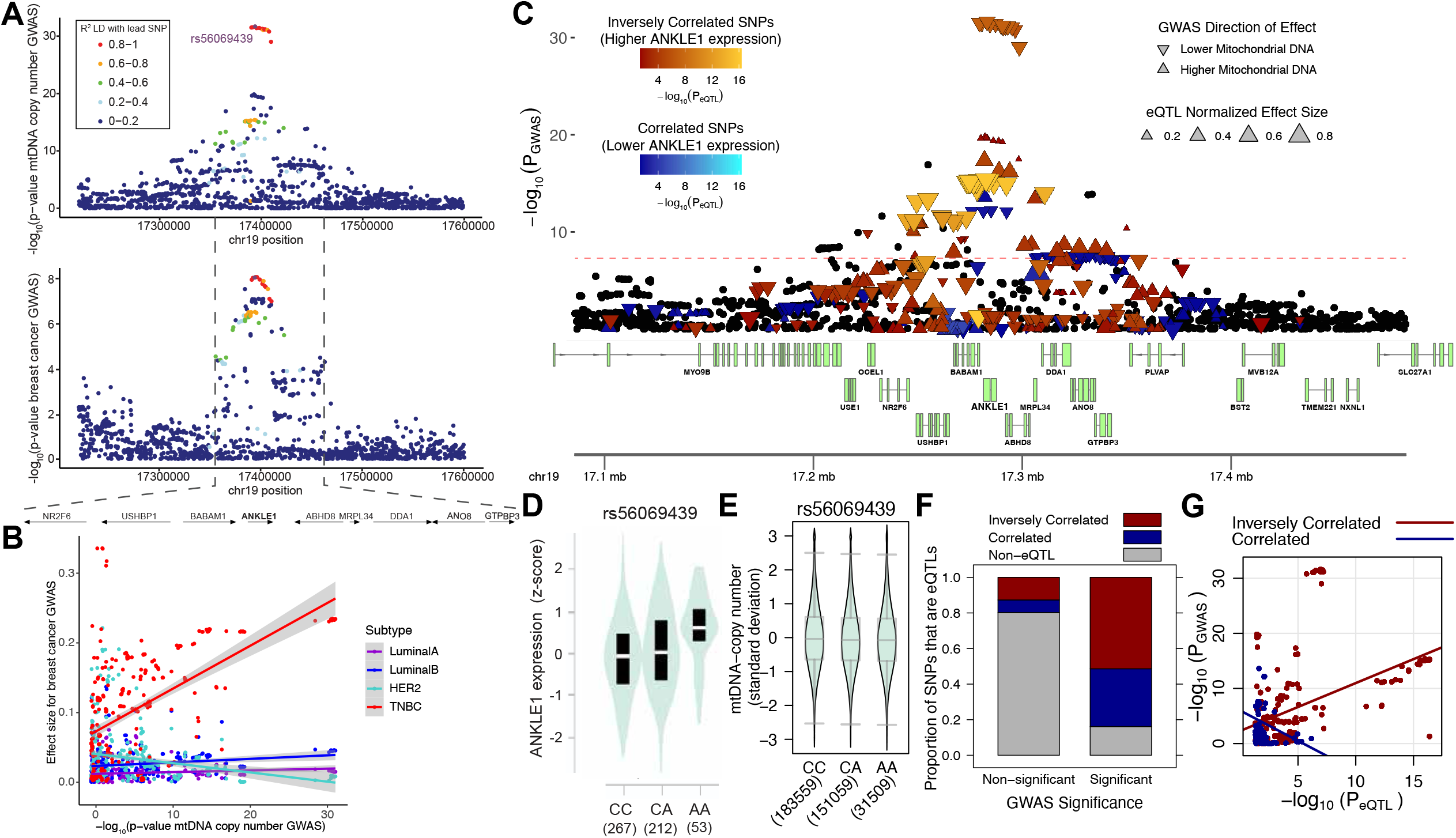
ANKLE1 expression inversely correlates with mitochondrial DNA copy number. A) Locus Zoom plots for mtDNA copy number and breast cancer risk show similar structure with a common linkage block for both phenotypes. Colocalization analysis indicates that mtDNA copy number and breast cancer risk likely share a single causal allele with 99% probability as measured by the coloc package (Wallace 2020). B) The relationship between breast cancer risk effect size and mtDNA-CN significance is specific to the TNBC breast cancer subtype. C) Integrative analysis of eQTL data from GTEx (Consortium et al. 2020) and mtDNA copy number GWAS (Chong et al. 2022) show that the GWAS alleles associated with lower mtDNA are associated with higher ANKLE1 expression (Drivas et al. 2021). D) The lead SNP for mtDNA copy number is an eQTL for ANKLE1. The minor allele associates with higher *ANKLE1* expression (Consortium et al. 2020). E) The minor rs56069439 allele associates with reduced mitochondrial DNA copy number (Chong et al. 2022). F) Over 50% of the genome-wide significant GWAS variants for mtDNA correlate with ANKLE1 eQTL in the expected direction (higher ANKLE1 associates with lower mtDNA). G) A scatter plot of log-transformed p-values from the mtDNA GWAS versus log-transformed pvalues from the ANKLE1 eQTL data shows evidence of colocalization.

Colocalization analysis indicates that the most significant mtDNA-CN GWAS variants are also eQTLs for *ANKLE1* (Figure 4C); moreover, the direction of effect is consistent with a role of ANKLE1 in digesting mitochondrial DNA. Higher *ANKLE1* expression alleles (Figure 4D) associate with lower mtDNA-CN (Figure 4E) (Chong et al. 2022). The majority of genome-wide significant alleles have an inverse correlation relationship between *ANKLE1* expression and mtDNA-CN (Figure 4F). The -log_10_ p-values for *ANKLE1* expression (eQTL) and mtDNA-CN (GWAS) are correlated exclusively for variants that display an inverse relationship between *ANKLE1* expression and mtDNACN (Figure 4G). Mitochondrial DNA is depleted in breast cancer tissue relative to normal tissue (Reznik et al. 2016).

These findings are consistent with the proposed role of AN-KLE1, which is to decrease the level of mtDNA in human cells. Taken together these integrative genomics analyses suggest that susceptibility to breast cancer, ANKLE1 expression, and mtDNA-CN phenotypes share common causal genetic variant(s) within the chr19p13.1 region.

### Expression of ANKLE1 leads to mitochondria degradation, DNA damage, Epithelial to Mesenchymal transition, and STAT1 activation

Since the molecular biology and genetic data indicate that ANKLE1 function is mediated through the mitochondria in normal and disease states, we hypothesized that ANKLE1 overexpression would lead to changes in expression of energy production and respiration genes. We performed genomic transcriptome profiling (RNA-seq) after overexpression of ANKLE1 for 1, 3, and 7 days (Figure S5A&C). Several gene set and gene ontology terms related to metabolic changes and mitochondrial regulation are enriched in the differentially expressed genes (Figure 5A&B and Figure S5D&E). *Hypoxia* and *glycolysis* are enriched gene sets among the activated genes (Figure 5A and Figure S5B). *Electron transport chain: OXPHOS system in mitochondria* is the most significant ontology term for repressed genes (Figure 5B).

**Fig. 5.**
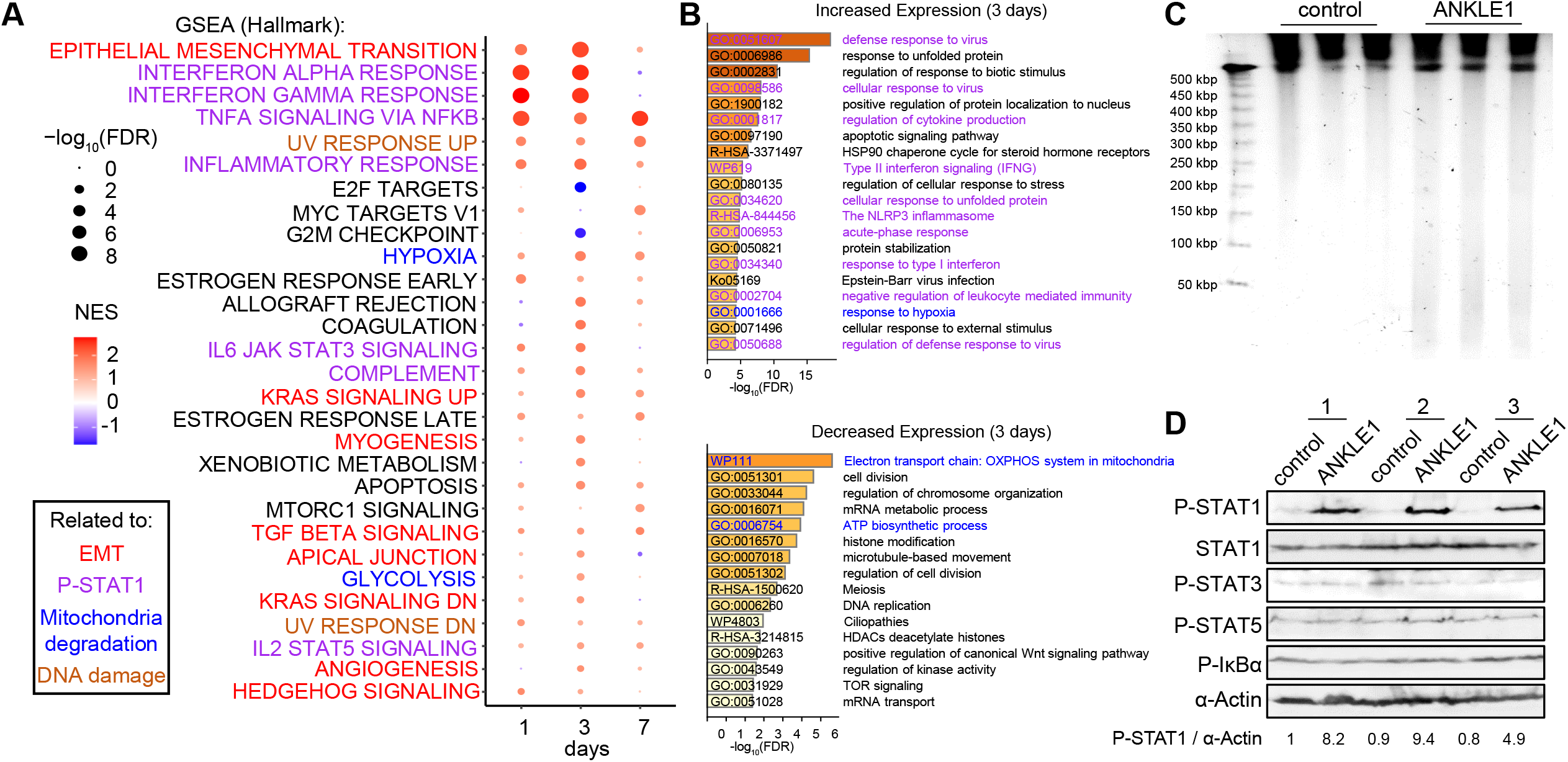
Expression of ANKLE1 leads to STAT1 activation, mitochondria degradation, DNA damage, and Epithelial to Mesenchymal transition. A) Gene Set Enrichment Analysis (GSEA) of RNA-seq results for ANKLE1 overexpressing cells versus control cells identifies hallmarks of STAT1 activation, mitochondria degradation, DNA damage, and Epithelial to Mesenchymal transition. B) Gene ontology analysis for differentially expressed genes (Figure S5C) after 3-days post-transfection of ANKLE1 cells versus control cells show enrichment for STAT1 activation and mitochondria degradation. C) ANKLE1 cuts nuclear DNA in non-apoptotic cells, as shown by DNA Pulse Field Gel Electrophoresis (PFGE) from non-apoptotic cells that overexpress ANKLE1 (24 hours post-transfection). D) ANKLE1 leads to STAT1 activation after 24 hours of ANKLE1-transfection, as shown by western blots probed for P-STAT1, STAT1, P-STAT3, P-STAT5, P-IκB and α-Actin of three independnet replicates per condition.

Previous work found that despite harboring endonuclease activity and both a nuclear import and export signal, ANKLE1 is not nuclearly localized and does not cause an extensive DNA-damage response (Brachner et al. 2012; Zlopasa et al. 2016). However, we observed that DNA damage response gene sets are activated upon ANKLE1 overexpression (Figure 5A and Figure S5B&D). We used a more sensitive assay, Pulse Field Gel Electrophoresis (PFGE), to assay ANKLE1’s ability to cut nuclear DNA when overexpressed. We excluded DNA breaks caused by apoptosis by selecting non-apoptotic cells (Figure S3A). We found that ANKLE1 cuts DNA with low frequency, resulting in fragments between 50 kbp and 200 kbp (Figure 5C).

Epithelial-mesenchymal transition (EMT) is the most significantly enriched gene set among ANKLE1 regulated genes (Figure 5A and Figure S5B). EMT is also a defining feature of breast cancer transformation (Felipe Lima et al. 2016; Micalizzi and Ford 2009). Several enriched gene sets represent hallmarks of EMT, such as TGF beta signaling, apical junction, and KRAS signaling (Figure 5A).

We found that many gene sets that relate to STAT1 signaling were enriched among the gene sets (Figure 5A&B and Figure S5B,D&E). Previous work found that mtDNA double strand breaks leads to phosphorylation of STAT1, which leads to genomic transcriptional changes (Tigano et al. 2021). Taken together with the mitochondria degradation phenotypes (Figure 3A&B), we hypothesized that mtDNA cleavage by ANKLE1 leads to STAT1 phosphorylation. Indeed, we found that STAT1 is phosphorylated in ANKLE1-expressing cells and this is not accompanied by activation of STAT3, STAT5, or NF-κB (Figure 5D). MtDNA damage synergizes with nuclear DNA damage to mount a robust type-I interferon response (Tigano et al. 2021) and the proteins DDX5 and cGAS are critical components of interferon signaling. cGAS is the major innate immune sensor of pathogenic DNA and can sense nuclear DNA breaks. DDX58 senses mtRNA exposed on the mitochondrial surface after mtDNA damage. We observe activation of DDX58 expression and inhibition of cGAS after ANKLE1 overexpression (Figure S5F). Taken together with the modest nuclear DNA damage after ANKLE1 overexpression (Figure 5C), this data is consistent with a mechanism by which mtDNA damage activates STAT1 upon ANKLE1 overexpression.

### ANKLE1 preferentially cleaves mitochondrial DNA

Variants in the chr19p13.1 locus both increase ANKLE1 expression and the risk of developing breast cancer. If ANKLE1 digests mtDNA in these cells, we expect that mtDNA content in breast cancer is inversely correlated with *ANKLE1* expression. We analyzed data that quantified mtDNA content and ANKLE1 expression in tumor samples and matched normal control samples (Reznik et al. 2016). Individuals with the highest ANKLE1 expression had consistently lower mtDNA content and mtDNA content is inversely correlated with *ANKLE1* expression when all patients are considered together (Figure S4A). We next sought to determine whether purified ANKLE1 (Figure S4B) prefers purified mitochondrial DNA as a substrate.

We isolated both mtDNA and nuclear DNA and incubated with ANKLE1 and monitored DNA degradation over 6 hours. ANKLE1 cleaves both DNA species, but mtDNA is degraded more rapidly and completely over the six hour time course (Figure 6A). Since previous work established that ANKLE1 preferentially cuts branched DNA (Song et al. 2020), we quantified the degradation of mtDNA with primers that span the D-loop and genic primers. The kinetics of degradation as measured by qPCR mirror the quantification by gel densitometry (Figure 6B). The D-loop contains brached DNA and consistent with our expectation, the D-loop mtDNA is degraded with faster kinetics and more completely than genic mtDNA (Figure 6B).

**Fig. 6.**
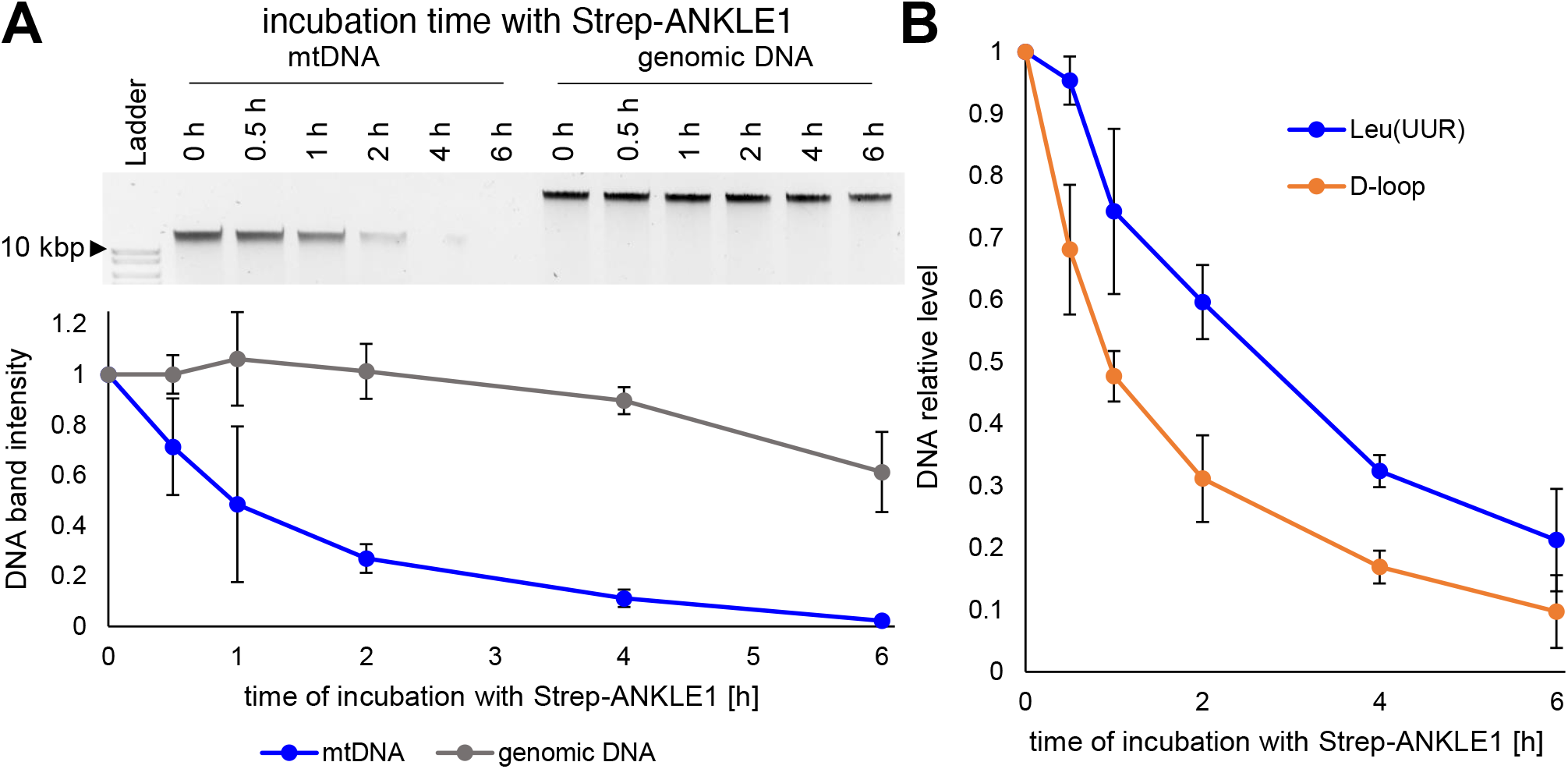
ANKLE1 preferentially cleaves mitochondrial DNA. A) The top gel is a representative gel of raw data of a time course treatment of purified mtDNA and nuclear DNA with purified ANKLE1. ANKLE1 degrades mtDNA faster than genomic DNA. The lower panel is quantification of three replicates. B) Quantitative PCR with primers flanking the D-loop or within the tRNA Leu(UUR) gene illustrates that ANKLE1 preferentially cuts the D-loop region of mtDNA. The standard deviation of three independent replicates is shown.

### ANKLE1 expression in normal breast epithelium cells drives the Warburg effect and apoptosis resistance in TP53 mutant cells

Recall that ANKLE1 expressing cells exhibit hallmarks of gene expression that indicate a switch from oxidative phosphorylation to glycolysis. In an effort to recapitulate the processes that may lead to transformation of normal breast cells, we began experiments in the MCF10A5E cell line. MCF10A-5E cells are clonally-derived from non-tumorigenic epithelial MCF10A cells (ATCC). These cells exhibit homogeneous behavior in 3D cultures and express many of the hallmarks of epithelial breast cells, such as epithelial sialomucins, cytokeratins, and milk fat globule antigen (Janes et al. 2010; Tait et al. 1990). Similarly to what we observed in HEK293T cells, ANKLE1 induces mitophagy in MCF10A-5E cells (Figure 7A). We found transient *ANKLE1* expression leads to chronic phenotypic changes within MCF10A-5E cells. We performed a 22 day time course experiment and we found that *ANKLE1* expression peaks within a day of transfection (Figure 7B). *ANKLE1* expression gradually decreases to background levels between 11-22 days after transfection (Figure S6A). mtDNA is decreased to approximately 50% and persists at this level for the remainder of the time course (Figure 7B). These data indicate that transient *ANKLE1* expression can lead to stable changes in cell metabolism.

**Fig. 7.**
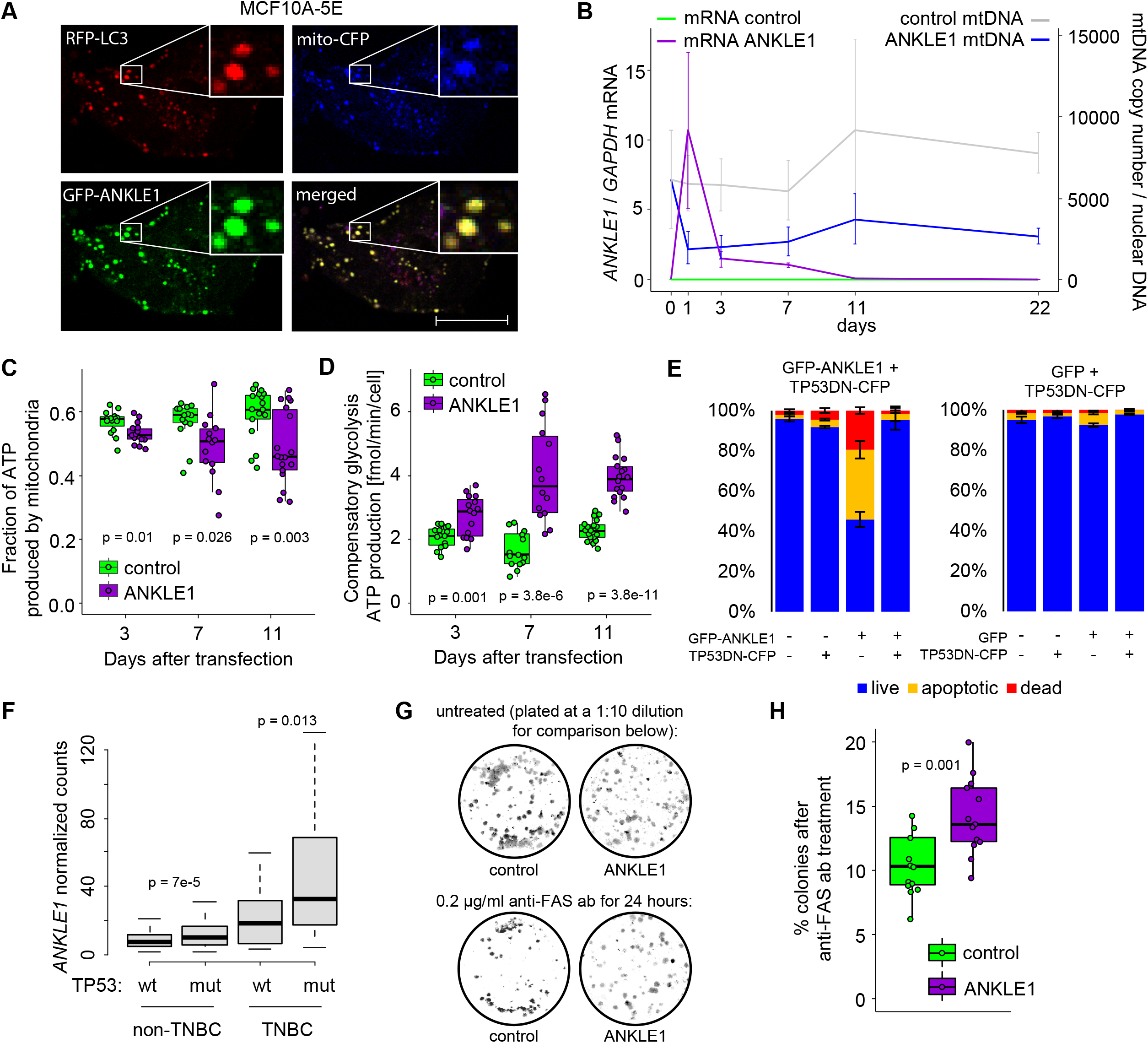
ANKLE1-induced mitophagy in normal breast cancer cells shifts the metabolism to glycolysis and increases resistance to apoptosis in TP53 negative cells. A) Confocal microscopy images (scale bar = 10 µm) of MCF10A-5E cells overexpressing GFP-ANKLE1, N_terminal_COX8-CFP, and RFP-LC3 proteins shows colocalization of ANKLE1, mitochondria, and autophagasomes. B) We quantified relative levels of *ANKLE1* mRNA and mtDNA over a 22 day time course of transient *ANKLE1* expression. Although *ANKLE1* levels return to baseline (Figure S6A), the decrease of mtDNA level is maintained. C-D) We used the Seahorse XF Real-Time ATP Rate Assay to show that ANKLE1 decreases the fraction of ATP produced by mitochondria (C) and ANKLE1 increases maximum compensatory glycolysis in MCF10A-5E (D). E) Flow cytometry analysis quantification of Figure S6E shows that ANKLE1 induces apoptosis in MCF10A-5E cells and apoptosis is abolished by a TP53 dominant negative mutant. F) The level of *ANKLE1* mRNA in human noTNBC and TNBC samples isolated from TP53 wild-type or mutant tumors indicates higher expression when TP53 is mutated. G-H) ANKLE1 overexpression produced more colonies when MCF10A-5E cells were treated with anti-FAS activating antibodies, even though ANKLE1 transient expression occurred 11 days prior to anti-FAS treatment (p-values are calculated with a two-tailed t-test).

We hypothesized that ANKLE1 expression contributes to a decrease in ATP produced by oxidative phosphorylation relative to ATP produced by glycolysis, which is known as the Warburg effect. We simultaneously measured extracellular acidification rate (ECAR) (Figure S6B) and oxygen consumption rate (OCR) (Figure S6C) using an ATP rate assay in ANKLE1-transfected and GFP-transfected control MCF10A-5E cells. We found higher acidification and lower oxygen consumption, which reflects lower oxidative phosphorylation and increased glycolysis. ECAR and OCR values were used to directly measure proton efflux rate (see Methods), which is an indirect measure of ATP production by mitochondria. We found that, on average, the fraction of ATP produced by the mitochondria is lower at days 3, 7, and 11 post-transfection (Figure 7C). Next, we blocked oxidative phosphorylation and quantified ATP produced by glycolysis in ANKLE1 vs. GFP control cells. We found that ANKLE1-transfected cells can produce ATP by glycolysis at a faster rate when oxidative phosphorylation is blocked, this is termed *compensatory glycolysis* (Figure 7D). We hypothesize that ANKLE1-mediated mitochondria degradation leads to a demand in energy resources that triggers activation of hypoxia (Figure 5A) and HIF1 pathways (Figure S5E), which lead to an increase in glycolysis. We repeated these experiments in HEK293T cells and we again found that ANKLE1 reduces the fraction of ATP produced in by the mitochondria and increases compensatory glycolysis (Figure S6D&E).

Throughout the experiments with ANKLE1 expressing MCF10A-5E cells, we empirically noticed that the fraction of apoptotic cells was higher than what we observed with ANKLE1 overexpression in HEK293T cells. The main genetic difference between these cell lines is the presence of large T antigen that deactivates TP53 in HEK293T cells. MCF10A-5E cells contain active TP53. Taken together with the fact that ANKLE1-mediated risk of breast cancer is primarily through its effect on the TNBC subtype, of which 80% are mutant for TP53 (Koboldt et al. 2012), we hypothesized that TP53 mutations allow cells that express ANKLE1 ectopically to evade apoptosis. We tested this hypothesis by transfecting MCF10A-5E cells with vectors carrying either GFP or GFP-ANKLE1 along with a CFP-TP53 dominant negative R175H mutant (TP53DN). We quantified apopto-sis after 24 hours with AnnexinV-PE and 7-AAD (Figure S6F). We found that ANKLE1-alone induces apoptosis in 60% of MCF10A-5E cells (Figure 7E). ANKLE1-driven apoptosis in MCF10A-5E cells is completely abolished in the presence of dominant negative TP53. In contrast, ANKLE1 does not increase apoptosis compared to the GFP control in HEK293T cells (Figure S6G). If mutant TP53 protects against ANKLE1-induced apoptosis, we would expect that TP53-mutant tumors have higher levels of ANKLE1 expression. We queried the Molecular Taxonomy of Breast Cancer International Consortium (METABRIC) database and we found that breast cancer tissues with mutated TP53 have significantly higher ANKLE1 expression irrespective of their TNBC status (Figure 7F). We experimentally tested the hypothesis that ANKLE1 overexpression will specifically induce apoptosis in TP53 wild type breast cancer cell lines. We compared ANKLE1-induced apoptosis in TP53 mutant breast cancer cells lines (HCC1806, MDA-MB-231 and MDA-MB-468) to TP53 wild type cells lines (MCF7, HCC1500 and CAL51). ANKLE1 increases apoptosis between 10% and 15% in p53-wild type cell lines and less than 3% in TP53 mutant cell lines (Figure S6H).

These support a model whereby ANKLE1 causes apoptosis in presence of wild-type TP53, and not in presence of mutant TP53. Mitochondria are known to play an essential role in apoptosis by regulating the release of proteins from the intermembrane space, which activates caspases (Wang and Youle 2009). To further highlight the importance of the mitochondria in apoptosis, previous work showed that mod-est differences in cellular mitochondrial content in clonal cell lines can determine the apoptotic fate upon activation of death receptor (Márquez-Jurado et al. 2018). We designed an experiment to directly test whether ANKLE1-induced depletion of the mitochondria (Figure 7B) increases the resistance of death receptor-triggered apoptosis in MCF10A5E cells. Recall that at day 11 ANKLE1 expression is at background levels, but mitochondrial DNA copy number is reduced by 50% (Figure S6A). We found that transiently expressed ANKLE1 prior to day 11 are more resistant to death receptor-induced apoptosis and able to proliferate and form colonies (Figure 7G&H).

### Transient, ectopic expression of ANKLE1 induces phenotypes consistent with hallmarks of carcinogenesis

Thus far we have shown that ANKLE1 promotes the Warburg effect and resistance to apoptosis within HEK293T and MCF10A-5E cells. Recall that the MCF10A-5E clone was selected for its ability to form homogenous 3D spheroids on the matrigel surface, which makes the model amenable for measuring features that are associated with transformation and cancer progression. MCF10A-5E cells cultured on matrigel form round hollow spheroids due to apoptosis of internally localized cells and this orderly growth is disrupted upon the introduction of an oncogene (Debnath et al. 2003; Janes et al. 2010). We transiently transfected ANKLE1 into MCF10A-5E cells and 11 days later we seeded them onto matrigel for an additional 11 days. We quantified the size and circularity of spheroids with OrganoSeg (Borten et al. 2018) (Figure S7A). Although MCF10A-5E cells no longer express ANKLE1 when they are seeded on matrigel, the spheroids that form are larger (Figure 8A) and less circular (Figure 8B and Figure S7B). We hypothesized that the spheroids that previously expressed ANKLE1 have a defect in apoptosis of internally localized cells, so we stained for cleaved Caspase 3 (a marker of apoptosis) and DNA (Figure 8C). Indeed, spheroids formed from acute ANKLE1expressing cells have more internally located intact nuclei (Figure 8D) and less apoptosis (Figure 8E). We confirmed less apoptosis in ANKLE1 spheroids by quantifying cleaved PARP protein levels by western blotting (Figure 8F). These data indicate that transient, ectopic expression of ANKLE1 in breast epithelium provides resistance to apoptosis, which can trigger spheroid phenotypes that are consistent with cancer progression.

**Fig. 8.**
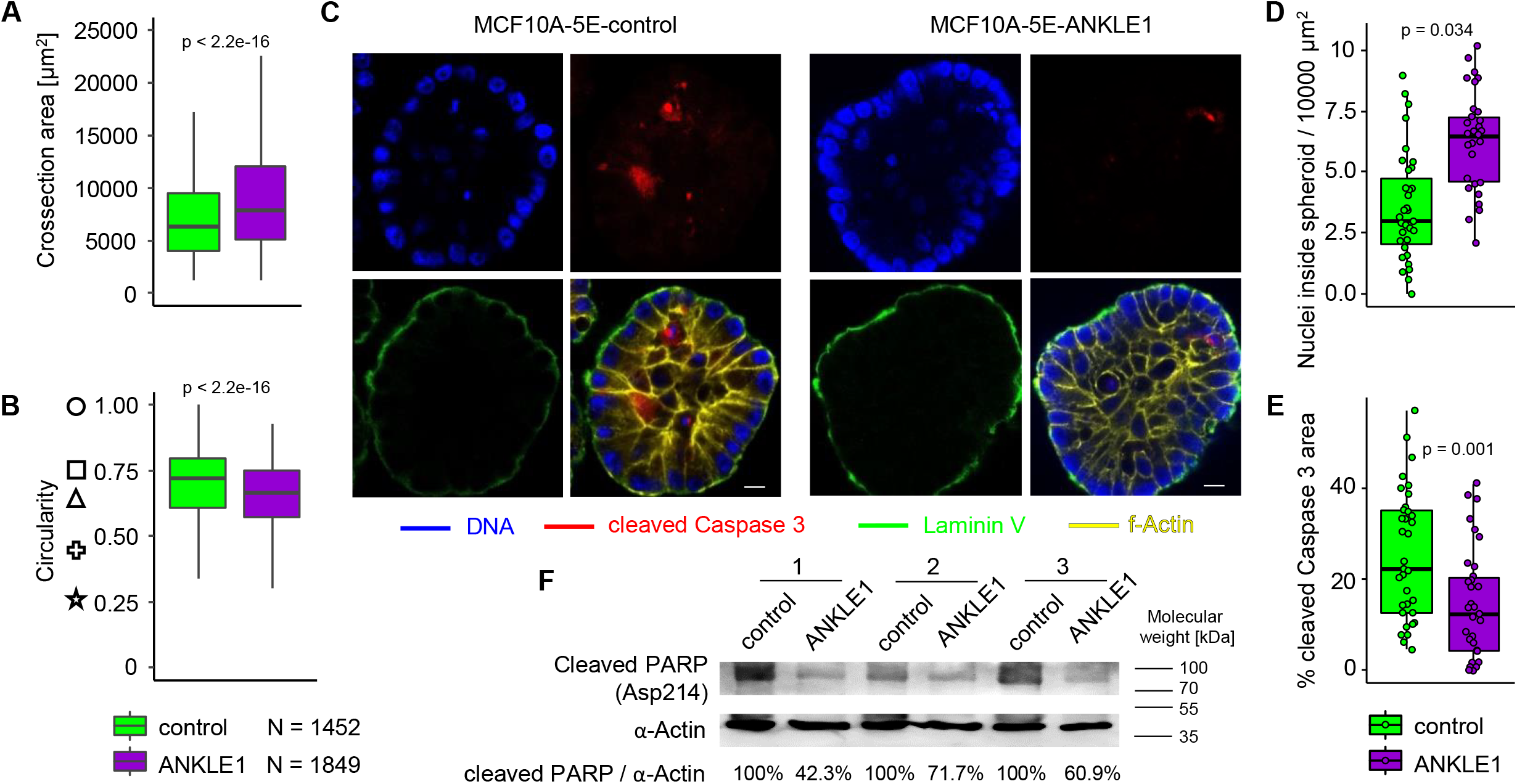
Spheroids formed by normal breast epithelium cells after ANKLE1 expression exhibit precancerous phenotypes. A) MCF10A-5E cells that transiently expressed ANKLE1 form larger spheroids. B) MCF10A-5E cells that transiently expressed ANKLE1 form less circular spheroids. Note that the symbols to the left are examples of circularity at the indicated y-value. C) Representative confocal microscopy images (scale bar = 10 µm) of spheroids stained for DNA, cleaved Caspase 3, and f-Actin reveal the effect of transient ANKLE1 expression. D-E) Spheroids derived from MCF10A-5E cells transiently transfected with ANKLE1 contain more intact nuclei (D) and exhibit less cleaved Caspase 3 staining (E) inside the spheroid. F) Western blot shows a lower level of cleaved PARP in protein extracts from spheroids derived from MCF10A-5E cells transiently expressing ANKLE1. We used densitometry to quantify the percent of cleaved PARP relative to the control.

## Discussion

Genome-wide association data and expression QTL data suggest common causal variant(s) within the chr19p13.1 locus for breast cancer risk, *ANKLE1* expression, and mtDNA copy number (Figure 9A). We propose that breast cancer risk variants directly regulate *ANKLE1* expression in breast tissue, which directly leads to mtDNA degradation and results in modified cellular metabolism and homeostasis.

**Fig. 9.**
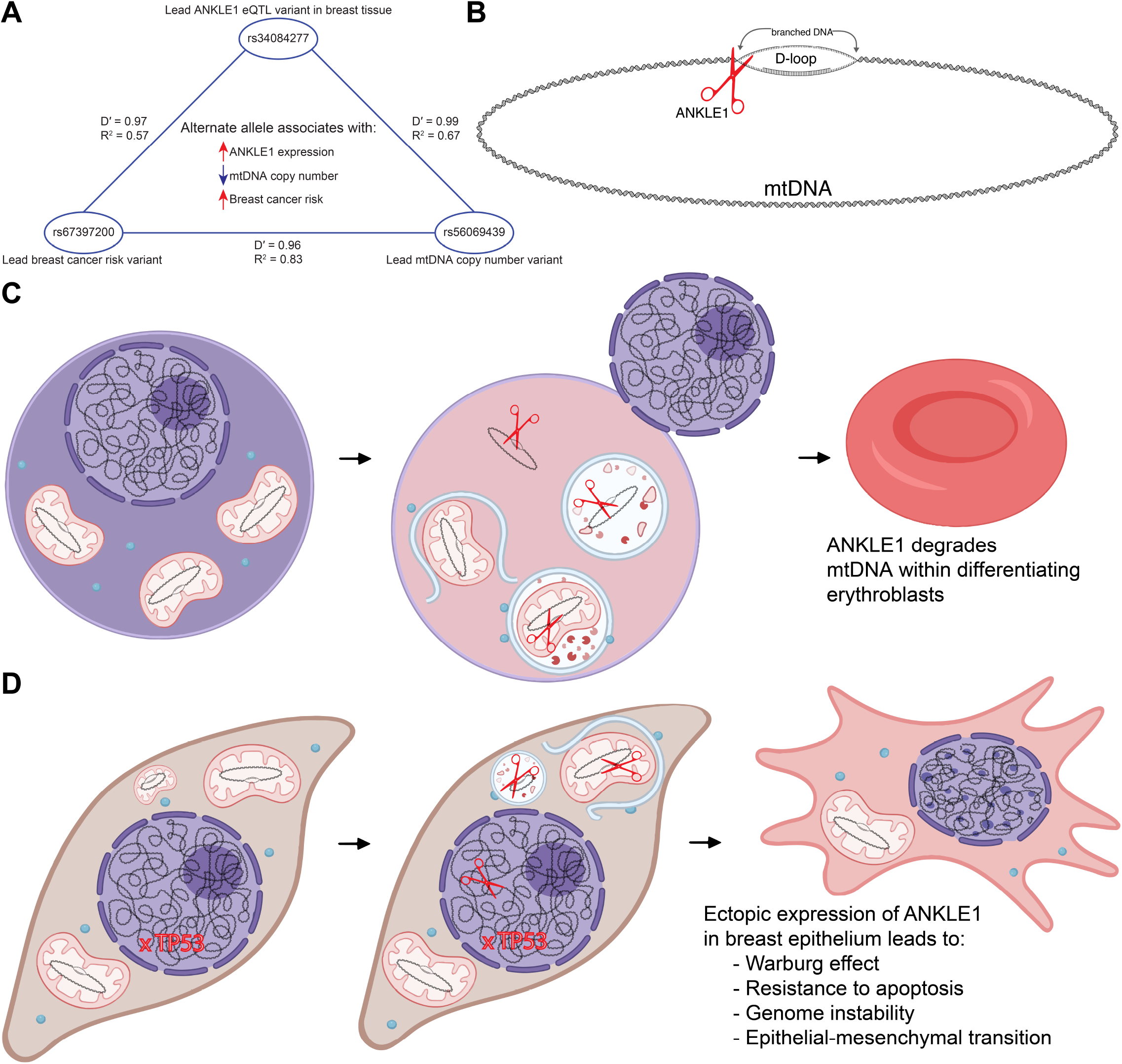
A model of ANKLE1 function in erythroblast differentiation and breast cancer. A) The lead genetic variants for breast cancer risk, *ANKLE1* expression, and mtDNA copy number are genetically linked to one another and each is associated with the other two phenotypes. B) ANKLE1 preferentially cuts branched DNA and we speculate that ANKLE1-mediated cleavage of the mtDNA D-loop facilitates degradation of mitochondrial DNA. C) We propose that the role of ANKLE1 in erythroblast differentiation is limited to mtDNA degradation. D) Ectopic expression of ANKLE1 in breast epithelium also causes mtDNA degradation, and this leads to mitophagy, the Warburg effect and resistance to apoptosis. Additionally, ANKLE1 in the nucleus may contribute to genome instability. Our RNA-seq data indicate that these phenotypes may lead to transcriptional changes that are consistent with the Epithelial-Mesenchymal transition.

Although there are relatively few reports on the function of ANKLE1, the data suggest that ANKLE1 originally evolved as a means to resolve DNA bridges (Hong et al. 2018a,b). Resolving DNA bridges is consistent with ANKLE1’s preferred specificity to cleave branched DNA (Song et al. 2020). While the molecular function of cleaving branched DNA remains unchanged in vertebrates, the ANKLE1 protein may have been functionally repurposed for mtDNA degradation in mammals. Mammals have evolved to degrade both their nuclei and mitochondria during erythropoiesis. The nucleus is actively expelled from the cell in a process known as enucleation and mitochondria are digested by mitophagy. Our data support a complementary mechanism to ensure degradation of the mtDNA in differentiating erythroblasts by ANKLE1 (Figure 2D and Figure S2H). The mitochondrial chromosome is circular and more stable because, without free DNA ends, circular DNA is refractory to digestion by exonucleases. Since mtDNA harbors a D-loop with branched DNA structures (Kasamatsu et al. 1971), *ANKLE1* cleavage of mtDNA may provide a mechanism to linearize the mitochodiral chromosome to ensure its faithful degradation (Figure 9B). ANKLE1mediated mtDNA digestion may prevent accumulation of cytosolic mtDNA that escaped from autophagosomes, as this phenomenon is associated with multiple metabolic diseases (Pérez-Treviño et al. 2020).

ANKLE1 is not expressed in normal breast tissue and we hypothesize that the ectopic expression of ANKLE1 leads to changes in metabolism, transcription, and cellular homeostasis. In support of this assertion, we show that ANKLE1 triggers cleavage of nuclear DNA in cells without triggering apoptosis (Figure 5C). The more prominent phenotype we observed is that ANKLE1 induces mtDNA degradation and mitophagy (Figure 3). The mitochondria phenotypes may drive STAT1 phosphorylation (Figure 5D) and a metabolic switch from oxidative phosphorylation to glycolysis (Figure 7).Consistent with these observations, we found modest DNA damage when ANKLE1 is overexpressed, but the most prominent phenotypes that we observed were mitochondrial phenotypes.

In the 1920s Otto Warburg showed that cultured tumor tissues have high rates of glucose uptake and lactate secretion, even in the presence of oxygen (aerobic glycolysis) (Warburg et al. 1927). Although cancer is caused by mutations within tumor suppressors and oncogenes, the Warburg effect represents a critical metabolic switch for cancer cells to survive and proliferate. There are several postulated roles of the Warburg effect in cancer cells. For instance a higher rate of ATP production by glycolysis may provide a selective advantage when competing for shared and limited energy resources. The Warburg effect has also been proposed to be an adaptation mechanism to support the biosynthetic requirements of uncontrolled proliferation. The increased glucose consumption is used as a carbon source for anabolic processes needed to support cell proliferation, such as the *de novo* generation of nucleotides, lipids, and proteins. As the tumor microenvironment has been more recently studied and appreciated to have a role in cancer biology, it has been proposed that the Warburg effect may present an advantage for cell growth in a relatively acidic and low oxygen microenvironment. Acidification of the microenvironment by lactate may allow for enhanced invasiveness and is a contributor to tumor-associated macrophage polarization that support tumor growth (Liberti and Locasale 2016). We suggest that ANKLE1-induced degradation of the mitochondria facilitates the Warburg effect to promote cancer growth.

ANKLE1-induced mitophagy may also promote survival of cancer cells by avoiding apoptosis. We show that ANKLE1 expression leads to lower mitochondria content and resistance to apoptosis caused by death receptor activation (Figure 7H&I). Activation of the death receptor is the primary mechanism by which the immune system removes dysregulated or mutated cells (O’Reilly et al. 2016). In support of this apoptosis-resistance hypothesis, previous work has shown that cells with fewer mitochondria are more resistant to apoptosis induced by death receptor activation (Márquez-Jurado et al. 2018). Another possible mechanism by which ectopic ANKLE1 expression may promote tumorigenesis in TP53 mutant cancers, which represent over 80% of TNBCs, is to increase mutational burden. We found that ANKLE1 cuts DNA in the nucleus without causing apoptosis in TP53 mutant cells, which will increase the mutation rate.

Modern genomics and phenotyping can convincingly identify genes that affect a phenotype, in this case ANKLE1 and breast cancer risk. A fundamental challenge is following up with experimental models that accurately recapitulate organismal biology. Alleles such as rs67397200, which af-fect ANKLE1 expression and contribute to breast cancer risk are common and found at 25% minor allele frequencies. Homozygosity for the risk allele (rs67397200) only affects the relative risk of developing breast cancer with an odds ratio of 1.17. Most GWAS hits have odds ratio in the range of 1.1-1.2. We concede that no model can accurately recapitulate the biology that occurs over 40 years of life that precedes a breast cancer diagnosis. Importantly, if we could recapitulate this biology note that the prevalence of breast cancer is low compared to allele frequency, so the vast majority of individuals with the risk allele do not develop breast cancer. A true model could be deemed a failure because the penetrance of disease incidence is too low. Moreover, ANKLE1 is one of hundreds of genes that contributes to risk of developing cancer, as opposed to a driver like p53, BRCA1, or Ras. Within the field of genetic epidemiology, following up on GWAS candidate genes to identify molecular mechanisms of risk is rate-limiting. With all these limitations in mind, the molecular phenotypes that we observe points to mitochondrial function. Although the system does not perfectly recapitulate the biology of living 40 years with slightly increased levels of ANKLE1 in breast epithelial tissue, the finding that ANKLE1-induces mitophagy in normal breast epithelium cells and shifts the metabolism to glycolysis, while increasing resistance to apoptosis in TP53 negative cells, is consistent with all our other data and models. Likewise, the fact that spheroids formed by normal breast epithelium cells after ANKLE1 expression exhibit precancerous phenotypes is supportive of the model that breast cancer risk attributed by ANKLE1 expression is mediated through ANKLE1’s effect on mitochondria biology.

In summary, the biological role of ANKLE1 is limited to degrading mtDNA in differentiating erythroblasts (Figure 9C). Ectopic ANKLE1 expression in normal breast cells leads to genome instability and degradation of mtDNA, which causes mitophagy, activation of STAT1, resistance to apoptosis, and a shift to aerobic glycolysis as a means of energy production (Figure 9D).

## Methods

### Analysis of publicly available data sets and statistical analysis

GWAS data were obtained from published metaanalysis summary statistics (Chong et al. 2022; Zhang et al. 2020). Expression QTL data were obtained from GTEx GCS bucket (https://console.cloud.google.com/storage/browser/gtexresources): dbGaP accession number phs000424.v8.p2 (Consortium et al. 2020). Violin plots of *ANKLE1* expression for different genotypes in breast tissue (Figure 1 and Figure 4) were obtained from the GTEx Portal. Colocalization analysis was performed with the coloc and eQTpLot packages (Drivas et al. 2021; Wallace 2020) using GWAS and QTL p-values to assess colocalization. Analyses we performed under the assumption that a single causal variant is responsible for both the eQTL phenotype and the GWAS phenotype. The effect sizes from the summary statistics were partitioned for different breast cancer subtypes and only the most significant (p<0.0001) GWAS single nucleotide polymorphisms (SNPs) were queried. *ANKLE1* expression in bone marrow subpopulations was analyzed based on microarray data: GDS3997, probe 1443978at (Konuma et al. 2011). *ANKLE1* expression during erythropoiesis was quantified by RNA-seq (GSE115684) (Ludwig et al. 2019). mtDNA copy number was previously measured by integrating whole exome and whole genome sequencing data (Reznik et al. 2016). Molecular Taxonomy of Breast Cancer International Consortium (METABRIC) data (Curtis et al. 2012) were accessed through the Xena platform (Goldman et al. 2020) and used to obtain TNBC, non-TNBC, normal adjacent, and TP53 mutation status, as well as *ANKLE1* expression (p-values were calculated with a two-tailed t-test). Circularity was calculated as: 4 π *Area / Perimeter ^2^. All the experiments were performed in at least three independent biological replicates.

### Plasmid construction

The pEGFP-C1-ANKLE1 plasmid that was used for all ANKLE1 expression experiments was kindly provided by Roland Foisner (Brachner et al. 2012). The pEGFP-C1 plasmid (Clontech V012024) was used as the control. sgRNA constructs for *ANKLE1* knockout were generated by inserting oligonucleotides containing the targeted sequences (5^*′*^-TTCAGGGCACAGCCTAGAAC -3^*′*^ and 5^*′*^-GATTCT-GCCCTAGCCCCACC -3^*′*^) into the pX458 vector (Addgene Plasmid #48138 (Ran et al. 2013)). Mito-CFP plasmid was obtained from Addgene (Plasmid #58426 (Mishra et al. 2014)). LC3-RFP plasmid was obtained from Addgene (Plasmid #21075 (Kimura et al. 2007)). TP53DN(R175H)-CFP was constructed by cloning CFP from mitoCFP plasmid into AgeI site of p53 (dominant negative R175H mutant)-pcw107-V5 (Addgene Plasmid #64638 (Martz et al. 2014)), using 5^*′*^-GGGTTAGGGATAGGCTTAC-CACCGGTTTACTTGTACAGCTCGTCCATGC -3^*′*^ and 5^*′*^-CTTGTACAAAGTGGTTACCGGAGGATC-CGGTGGTGTGAGCAAGGGCGAGGAGCTG -3^*′*^ primers for PCR and In-Fusion Cloning (Takara Bio). The pStrep-ANKLE1 plasmid was obtained by cloning annealed oligonucleotides containing Twin-Strep-tag (5^*′*^ CCGGTCACCATGGCGTGGAGCCACCCGCAGTT-CGAGAAAGGTGGAGGTTCCGGAGGTGGATCGG-GAGGTTCGGCGTGGAGCCACCCGC-AGTTCGAAAAAGC 3^*′*^ and 5^*′*^ GGCCGCTTTTTCGAACTGC-GGGTGGCTCCACGCCGAACCTCCCGAT-CCACCTCCGGAACCTCCACCTTTCTCGAA-CTGCGGGTGGCTCCACGCCATGGTGA 3^*′*^) into AgeI, NotI fragment of pEGFP-C1-ANKLE1 to remove GFP.

### Cell culture, treatment, transfection, and generation of stable cell lines

K562, MCF7, CAL-51, HCC1500, MDA-MB-468, HCC1806, MDA-MB-231 and HEK293T cells were purchased from ATCC. MCF10A-5E cells were kindly provided by Kevin Janes. K562 cells were cultured in RPMI supplemented with 10% FBS (growth condition) or in RPMI (-glutamine) supplemented with 10% FBS and 1mM sodium butyrate (differentiation medium) (Canh Hiep et al. 2012). MCF7, CAL-51, MDA-MB-468, MDA-MB-231 and HEK293T cells were cultured in DMEM supplemented with 10% FBS. HCC1500 and HCC1806 cells were cultured in RPMI supplemented with 10% FBS. MCF10A-5E cells were cultured in DMEM/F12 supplemented with 5% horse serum, EGF (20 ng/ml), hydrocortisone (0.5 µg/ml), cholera toxin (0.1 µg/ml stock), and insulin (10 µg/ml) (Debnath et al. 2003). Cells were first transient transfected and cultured for 11 days, then the cells were seeded on the surface of matrigel (Thermo Fisher Scientific) in DMEM/F12 supplemented with 2% horse serum, EGF (2 ng/ml), hydrocortisone (0.5 µg/ml), cholera toxin (0.1 µg/ml stock), insulin (10 µg/ml), and 2% matrigel for another 11 days to form spheroids (Debnath et al. 2003). Cells were treated with MitoTracker™ Red CMXRos (100 nM) and LysoTracker™ Deep Red (50 nM) 30 min before fixation or live imaging. Bafilomycin A1 (InvivoGen) was added to cells (10 nM final concentration) 6 hours after transfection, flow cemetery was performed 18 hours later (24 hours after transfection). All cell lines were transfected with Lipofectamine 3000 reagent. K562 *ANKLE1* KO lines were obtained by co-transfecting two plasmids carrying sgRNAs for *ANKLE1* or by transfecting pX458 as a control. The GFP-positive cells were sorted into 96-well plates one day after transfection. Clones were validated by PCR with the following primers: 5^*′*^GGTTAGTCTTCCCAGGGCAC -3^*′*^ and 5^*′*^GCCTCCCGTGTATAAGCCTC -3^*′*^ (1838 bp PCR product for WT clones vs. 270 bp PCR product for KO clones). Hemoglobin was measured by absorbance at 425 nm of 10^5^ cells suspended in 0.1 ml of PBS. For the colony formation assay, cells were treated for 24 h with 500 ng/ml Anti-Fas (human, activating) clone CH11 antibody (Milipore 05-201) and seeded on 6-well plates at the specified dilutions. In the top panel of Figure 7G, only 10% of untreated cells were imaged to ensure that the number of colonies were visually comparable to the antibody-treated cells. Colonies were fixed with methanol and stained with Enhanced Gram Crystal Violet Ethanol Solution (ThermoFisher Scientific).

### Strep-ANKLE1 purification and DNA digestion

We transfected pStrep-ANKLE1 plasmid into HEK293T cells for 24 hours and performed affinity purification MagStrep3XTBeads and BXT elution buffer (IBA Lifesciences) according to the manufacturer’s protocol. Genomic DNA and mtDNA were isolated from HEK293T cells with DNeasy Blood & Tissue Kit (Qiagen) and Mitochondrial DNA Isolation Kit (abcam). 1*µ*g of either mitochondrial or nuclear DNA was incubated with 100 ng of Strep-ANKLE1 in a buffer containing: 20 mM HEPES-KOH pH 7.4, 2 mM MnCl2, 45 mM KCl, 50 mg/ml BSA. DNA was purified by ethanol precipitation for qPCR or directly run on 0.8% agarose gel.

### qPCR

Genomic DNA was isolated with QuickExtract™ DNA Extraction Solution. RNA isolation, cDNA synthesis, and qPCR were performed with Power SYBR™ Green Cellsto-CT™ Kit using StepOne™ Real-Time PCR System (Applied Biosystems). Mitochondria content was measured by primers specific for human mtDNA Leu(UUR) region (primers: 5^*′*^-CACCCAAGAACAGGGTTTGT -3^*′*^ and 5^*′*^-TGGCCATGGGTATGTTGTTA -3^*′*^) and genomic DNA (primers: 5^*′*^-GAGGCAGGACTCAGGACAAG -3^*′*^ and 5^*′*^-GGATGCCTCAGGGACCAG -3^*′*^). *ANKLE1* RNA level was measured by primers specific for *ANKLE1* (primers: 5^*′*^-ACACCCTTCACCAGGCAGTT -3^*′*^ and 5^*′*^-AAAACTCTGGGCCAGGAGCAA -3^*′*^) and we used the following *GAPDH* primers: 5^*′*^-TGCACCACCAACTGCT-TAGC -3^*′*^ and 5^*′*^-GGCATGGACTGTGGTCATGAG -3^*′*^.

### Immunofluorescence and microscopy

Cells or spheroids were fixed with 4% PFA (20% Paraformaldehyde Solution (Electron Microscopy Science)) for 1 h. Spheroids were blocked and permabilized with 0.5 % Triton, 100mM Glycine, 10 % FBS in PBS for 30 min, then stained overnight with Laminin-5 Alexa Fluor 488 Conjugated antibodies (Milipore), Cleaved Caspase-3 (Asp175) (D3E9) Alexa Fluor 467 antibodies (Cell Signaling) and ActinRed 555 (ReadyProbes, Invitrogen) overnight in 0.2 % Triton, 10 % FPS in PBS. Stained spheroids were mounted in ProLong Gold antifade reagent with DAPI (Invitrogen) on glass coverslips and imaged with Zeiss LSM 710 Multiphoton confocal microscope. Images were quantified using the Fiji software (Schindelin et al. 2012).

### Flow cytometry

Enucleation was measured by staining with DRAQ5 (Fisher Scientific) and Phalloidin-FITC (TOCRIS) and assessing the fraction of DRAQ5 negative cells. Apoptosis was detected by staining with 7-AAD and Anexin V-PE (Apoptosis Detection Kit I - BD Pharmingen). Attune NxT flow cytometer (Life Technologies) was used for all flow cytometry experiments. For imaging flow cytometry live cells were stained with DAPI (Sigma) and run on Amnis ImageStreamX Mark II (Luminex). Cells were sorted with Influx Cell Sorter (BD Biosciences). Flow cytometry data were analysed with FCS express software.

### Western blot

Cells were lysed in IPH buffer (50 mM Tris-Cl, 0.5% NP-40%, 50 mM EDTA). Protein lysates were run on 10% polyacrylamide SDS-PAGE gels and transferred to nitrocellulose membranes (Amersham Protran 0.45 µm NC - GE Healthcare Life Sciences). Membranes were blocked for 30 minutes in 5% milk in TBST buffer and incubated overnight with primary antibody (1:1000). Secondary antibody (1:5000) incubation was carried out for 1 hour after washing with TBST, and before washing and incubation with SuperSignal™ West Pico PLUS Chemiluminescent Substrate (ThermoFisher). The following primary antibodies were used: Anti-Phospho-Stat1 (Tyr701) (58D6) antibody (Cell Signaling Technology 14994), Anti-Stat1 (D1K9Y) antibody (Cell Signaling Technology 80916), Anti-Phospho-Stat3 (Tyr705) (D3A7) antibody (Cell Signaling Technology 9145), Anti-Phospho-Stat5 (Tyr694) (D47E7) antibody (Cell Signaling Technology 4322), AntiPhospho-IκBα (Ser32) (14D4) antibody (Cell Signaling 2859), Anti-Cleaved PARP (Asp214) (D64E10) antibody (Cell Signaling Technology 5625) and Anti-β-actin (AC-74) antibody (GenWay Biotech Inc. GWB-A0AC74). Densitometry was performed with Fiji(Schindelin et al. 2012).

### ATP rate assay

Percent ATP produced by mitochondria and the level of compensatory glycolysis were measured using a Seahorse XF Real-Time ATP Rate Assay Kit and XFe96 Analyzers (Agilent) according to manufacturer instructions. Seahorse XF Analyzers directly measure real time extracellular acidification rate (ECAR) and oxygen consumption rate (OCR) of cells. ECAR and OCR are indicators of the two major energy-producing pathways: glycolysis and oxidative phosphorylation. Basal OCR and ECAR rates were first measured. Injection of oligomycin results in an inhibition of mitochondrial ATP synthesis that results in a decrease in OCR, allowing for quantification of mitochondrial ATP Production Rate. ECAR data combined with the buffer factor of the assay medium allows calculation of total Proton Efflux Rate (PER) using the following formula: PER (pmol H+/min) = ECAR (mpH/min) × BF (mmol/L/pH) × Geometric Volume (µL) × Kvol. Complete inhibition of mitochondrial respiration with rotenone plus antimycin A accounts for mitochondrial-associated acidification, and when combined with PER data allows calculation of the glycolytic ATP Production Rate. Compensatory glycolysis was measured as ATP production with complete inhibition of mitochondrial respiration.

### Pulse Field Gel Electrophoresis (PFGE)

Live cells were embedded in 1% LMP agarose in 50mM EDTA (pH 8). Solidified plugs were incubated in 1% LDS (RPI Research Products International), 100 mM EDTA, 10 mM Tris pH 8.0 overnight and then washed 5 times (30 min each wash) in 50 mM EDTA (pH 8). Plugs were then secured in 1% Certified Megabase Agarose (BIO-RAD) gel in 0.5x TBE buffer and run with Lambda Ladder (ProMega-Markers) using CHEF-MAPPER system (BIO-RAD).

### RNA-seq

Live, nonapoptotic, GFP-positive cells were sorted 1, 3 and 7 days after transfection with pEGFP-C1-ANKLE1 or pEGFP-C1 plasmids. RNA was isolated by TRIzol extraction using Direct-zol RNA MicroPrep Kit including DNase treatment (ZymoResearch). rRNA depletion was performed using a NEBNext rRNA Depletion Kit v2 (Human/Mouse/Rat) and RNA was purified using Agencourt RNAClean XP Beads (New England Biolabs). Libraries were prepared using the NEBNext Ultra II Directional RNA Library Prep Kit for Illumina (New England Biolabs). Concentrations of libraries were measured with Qubit dsDNA HS Assay Kit (Thermo Fisher Scientific) and fragment size distribution were measured with TapeStation (Agilent). Libraries were pooled together and sequenced at the University of Virginia Genome Analysis and Technology Core using the Illumina NextSeq2000 instrument. RNA-seq data were aligned to the human assembly hg38 (Gencode v33) using HISAT2 (Kim et al. 2015) and quantified by HTSeq (Anders et al. 2015). We applied DESeq2 (Love et al. 2014) to identify differentially expressed genes with a false discovery rate less than 0.05 (Figure S5C). Principal component analysis was performed to ensure that replicates group together and that variation is observed between control and ANKLE1 expressing cells and time points, rather than between batches (Figure S5A). Gene Set Enrichment Analysis (GSEA) (Subramanian et al. 2005) was performed in R with the fgsea package (Korotkevich et al. 2021) using log_2_(Fold Change)*-log_10_(FDR) to rank all genes. Gene ontology analysis was done using Metascape (Zhou et al. 2019) on significantly (FDR < 0.05) activated or repressed genes. All RNA-seq library data files are available under GEO accession number GSE186393.

## ACKNOWLEDGEMENTS

This work was funded by R35-GM128635 to MJG. RP was supported by an NCI Predoctoral to Postdoctoral Fellow Transition Award (F99/K00-CA253732). We want to thank University of Virginia Flow Cytometry Core (RRID: SCR_017829), University of Virginia Genome Analysis and Technology Core (RRID: SCR_018883) and University of Virginia Advanced Microscopy Facility (RRID: SCR_018736).

## AUTHOR CONTRIBUTIONS

PP and MJG conceptualized and developed the project. PP performed all experiments with the exception of the RNA-seq experiment performed by RP. PP and MJG analyzed all the data and wrote the manuscript.

---

**Fig. S1.**
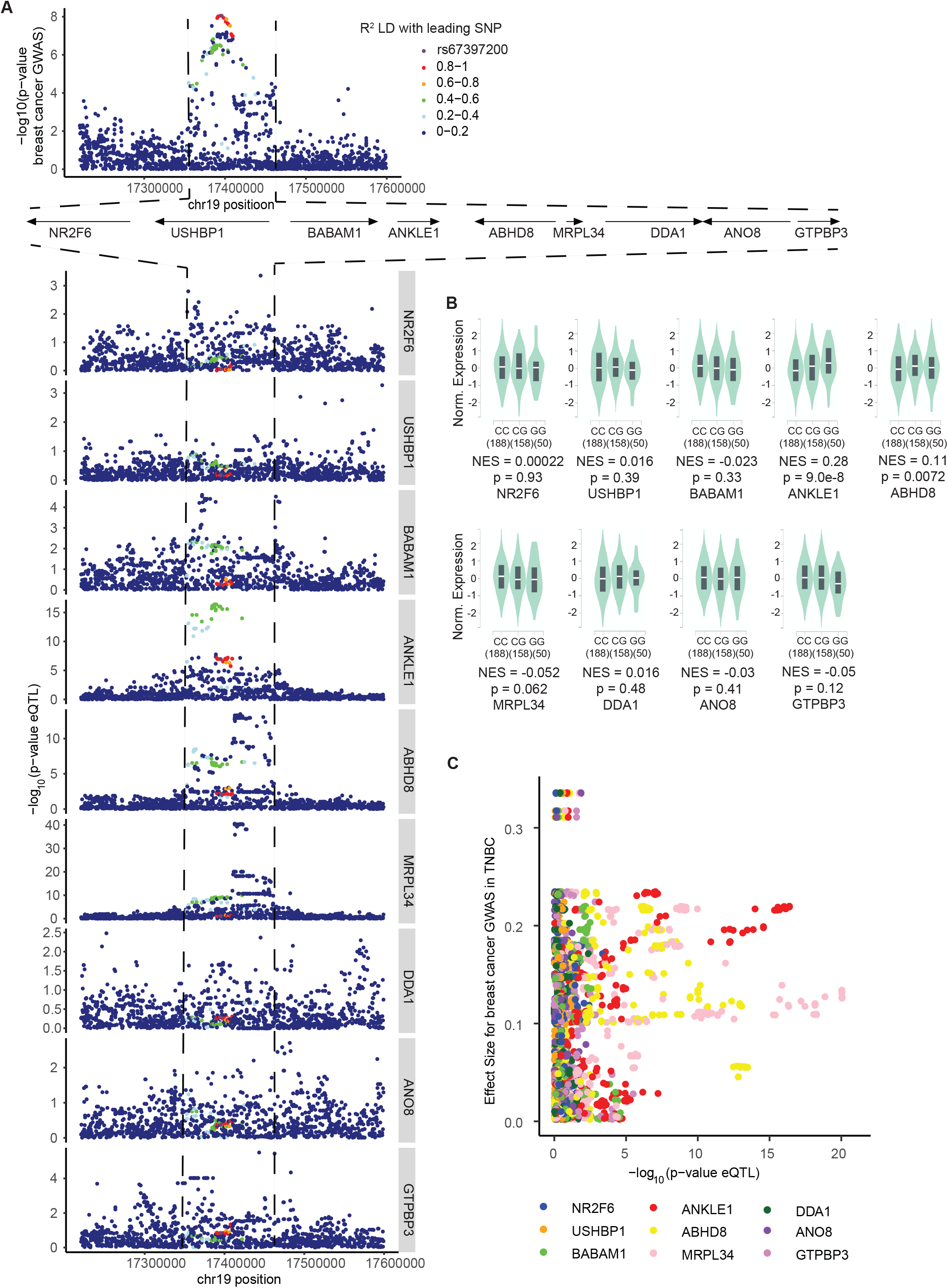
Comparison of *ANKLE1* with other genes in the locus confirms that *ANKLE1* is the most likely causal breast cancer susceptibility gene within chr19p13.1. A) The breast cancer susceptibility GWAS variants within the chr19p13.1 locus (higher panel) colocalize predominantly with *ANKLE1* eQTL variants as opposed to the other local genes (lower panel). B) The G allele of rs67397200, which is associated with increased breast cancer risk, is associated with higher expression of *ANKLE1* in breast tissue. The risk allele does not associate with the expression of other genes in the locus, except *ABHD8* albeit with a less significant p-value. C) The *ANKLE1* eQTL p-value correlates best with TNBC GWAS effect size.

**Fig. S2.**
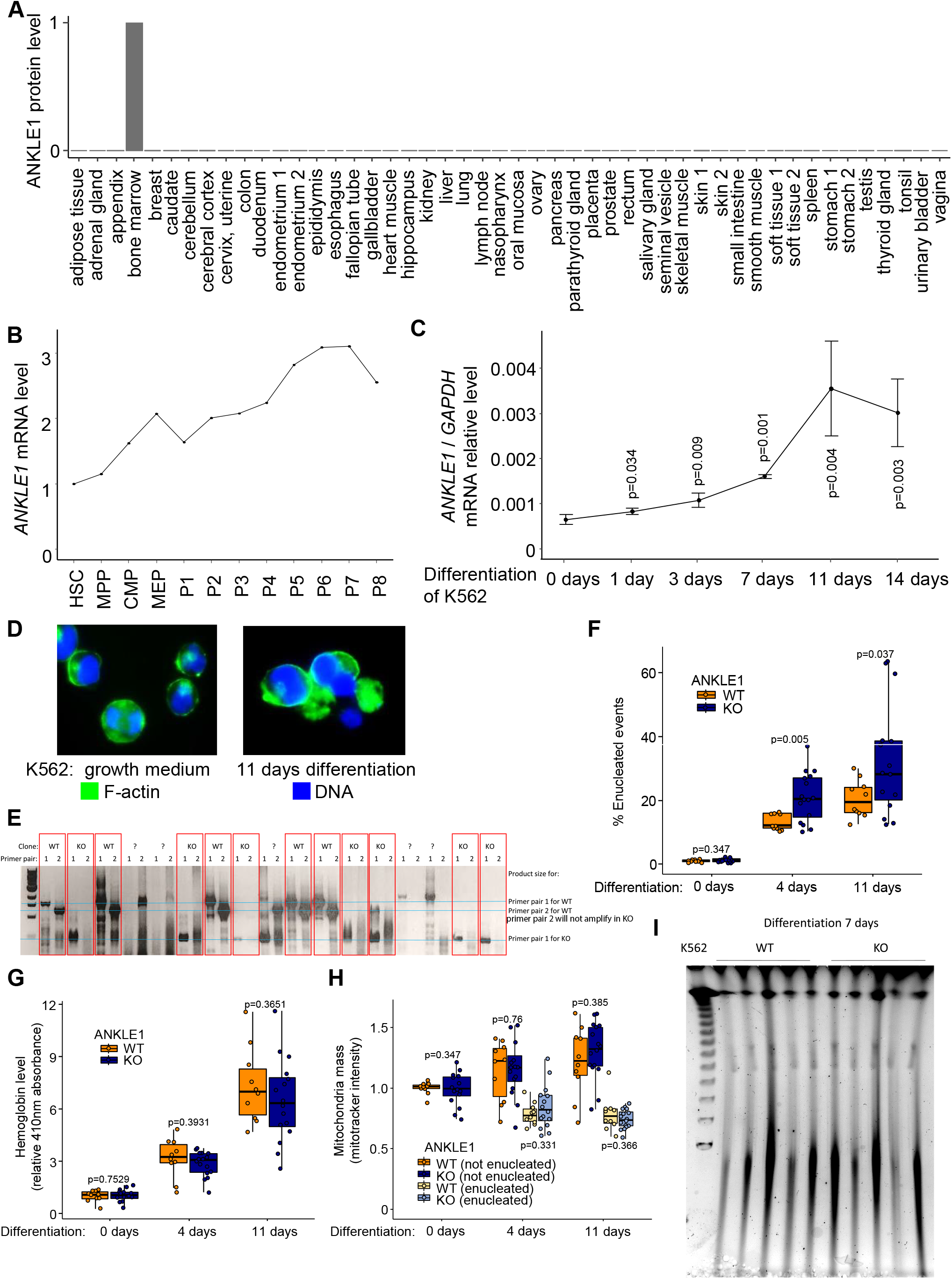
*ANKLE1* expression is only within the bone marrow and increases throughout erythropoiesis, but knockout of *ANKLE1* has only modest phenotypes. A) ANKLE1 protein expression in human tissues is limited to bone marrow (https://www.proteinatlas.org/). B) *ANKLE1* mRNA level increases throughout erythroid differentiation (RNA-seq data: GSE115684 (Ludwig et al. 2019)): hematopoietic stem cells (HSC), multipotential progenitors (MPP), common myeloid progenitors (CMP), megakaryocyte erythroid progenitors (MEP), myeloid progenitor (P1) to reticulocyte (P8). C) *ANKLE1* mRNA increases in differentiating K562 cells. P-values result from a T-test compared to time point 0. D) A representative picture of K562 cells shows enucleation after 11 days in differentiation medium (right panel) compared to undifferentiated cells (left panel). E) PCR validates the K562 knockout clones and tehir wild type counterparts. F) The fraction of enucleated cells is increased in *ANKLE1* KO K562 cells during differentiation. G) Hemoglobin level is unchanged upon *ANKLE1* KO in K562 cells during differentiation. H) *ANKLE1* KO does not affect mitochondria mass during K562 cell differentiation (p-values are calculated with a two-tailed t-test). I) Nuclear DNA fragmentation is no different between wild-type (WT) and knockout ANKLE1 K562 cells, as shown by pulse field gel electrophoresis analysis of DNA isolated from undifferentiated and differentiated (7 days) K562 cells.

**Fig. S3.**
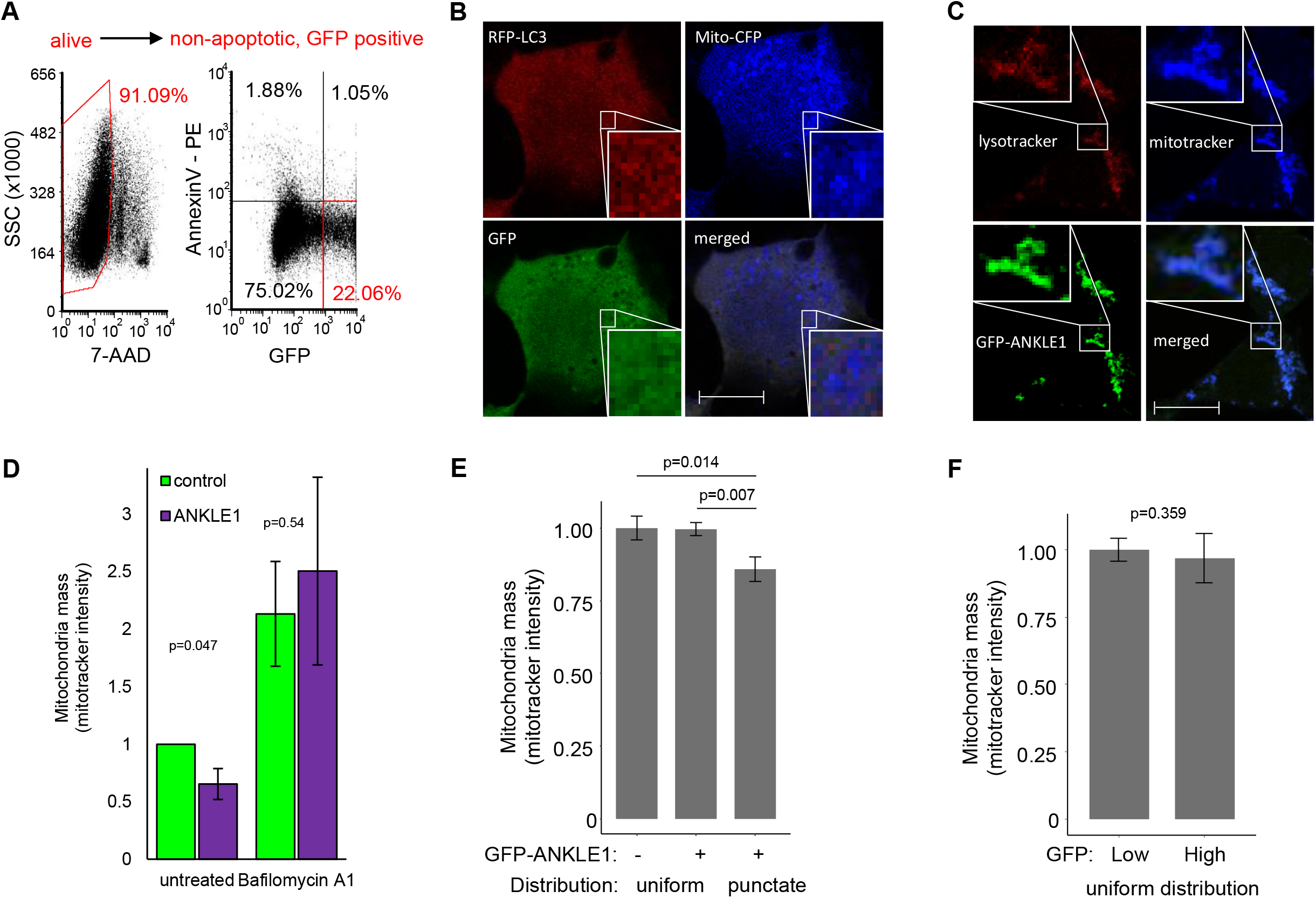
*ANKLE1* expression correlates with decreased mtDNA and ANKLE1 protein localizes to the mitochondria. B) *ANKLE1* expression negatively correlates with mtDNA fold change between tumor and normal tissue in breast cancer (Reznik et al. 2016). A) Alive, non-apoptotic and GFP-positive cells were selected using the illustrated gating strategy. B) Confocal microscopy imaging (scale bar 10 µm) with fluorescent-tagged mitochondrial targeting sequence present in the N-terminus of COX8, GFP, and LC3 proteins shows uniform distribution of GFP and lack of colacalizatino between GFP, LC3, and mitochondria. This control is quantified within Figure 3C. C) Confocal microscopy images (scale bar 10 µm) of HEK293T cells overexpressing GFP-ANKLE1, stained with mitotracker and lysotracker show colocalization of ANKLE1, mitochondria, and lysosomes, as an orthogonal method to Figure 3C. D) Bafilomycin treatment abrogates the effect of ANKLE1 over-expression reducing mitochondrial mass. E) Imaging flow cytometry of HEK293T cells overexpressing GFP-ANKLE1 indicates that ANKLE1 expression causes lower mitochondria content (measured by mitotracker fluorescence) only when ANKLE1 is distributed in puncta. F) In contrast, differences in ANKLE1 intensity does not cause lower mitochondria content when ANKLE1 is uniformly distributed throughout the cell (p-values are calculated with a two-tailed t-test).

**Fig. S4.**
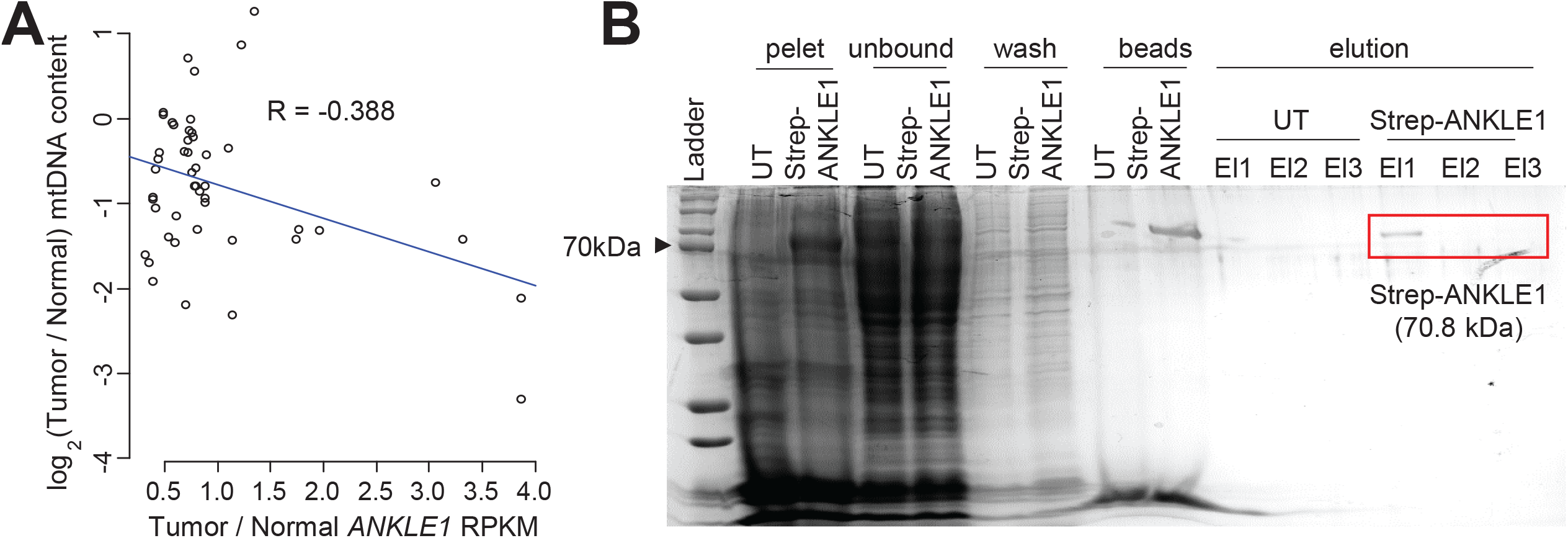
*ANKLE1* expression inversely correlates with mtDNA fold change between tumor and normal tissue in breast cancer. A) The log ratio of Tumor/Normal mtDNA content on the y-axis is plotted against the log ratio of tumor/normal ANKLE1 expression on the x-axis (Reznik et al. 2016). B) We purified the ANKLE1 protein by Streptavidin-tag affinity chromatography. The UT lanes corresponds to untransfected HEK293T cell lysate purification and Strep-ANKLE1 lanes correspond to purification from HEK293T cells transfected with pStrep-ANKLE1 vector. Purified Strep-ANKLE1 is 70.8 kDa.

**Fig. S5.**
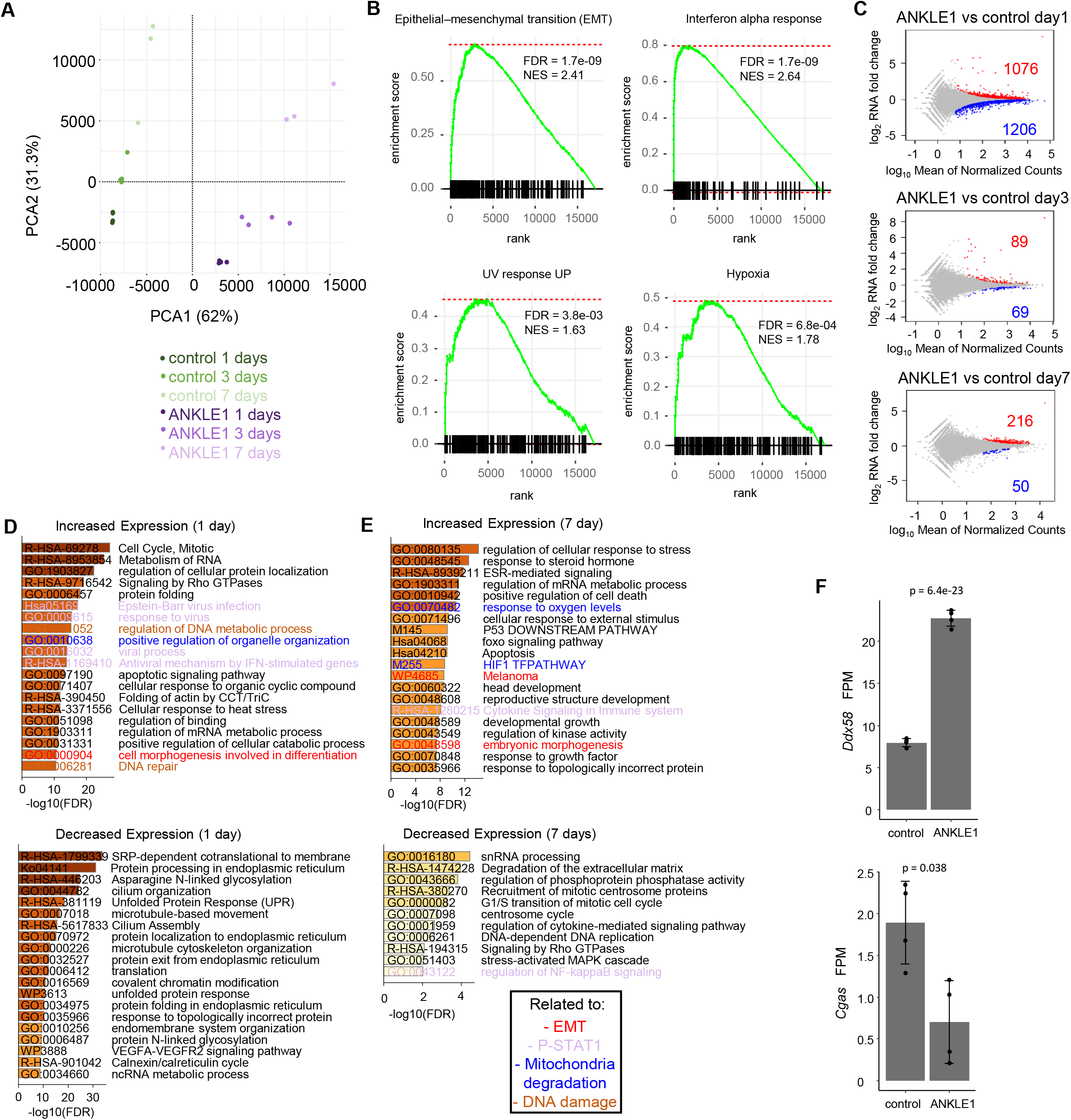
RNA-seq analysis of all time points indicates that ANKLE1 expression leads to changes in EMT, STAT1 activation, mitochondria degradation, and DNA damage. A) Principal component analysis of the RNA-seq data illustrates that biological replicates cluster together. B) GSEA plots show enrichment of genes engaged in Epithelial-Mesenchymal transition, Interferon alpha response, UV response UP, and Hypoxia. C) These plots compare the change in expression of all genes upon *ANKLE1* transfection for the indicated times. Red genes are activated upon ANKLE1 expression, blue genes are repressed, and light-gray points represent all other genes. The numbers of activated and repressed genes are indicated by text with the corresponding color. D-E) Gene ontology analysis for differentially expressed genes (Figure S5C) for the 3 day (D) and 7 day (E) time points after transfection. ANKLE1 overexpressing cells versus control cells show enrichment for STAT1 activation DNA damage and mitochondria degradation. F) DDX58 senses mtRNA exposed on the mitochondrial surface after mtDNA damage. cGAS is the major innate immune sensor of pathogenic DNA and can sense nuclear DNA breaks. We observe activation of DDX58 expression and inhibition of cGAS after ANKLE1 overexpression. FDR adjusted p-values from the DESeq2 analysis are shown.

**Fig. S6.**
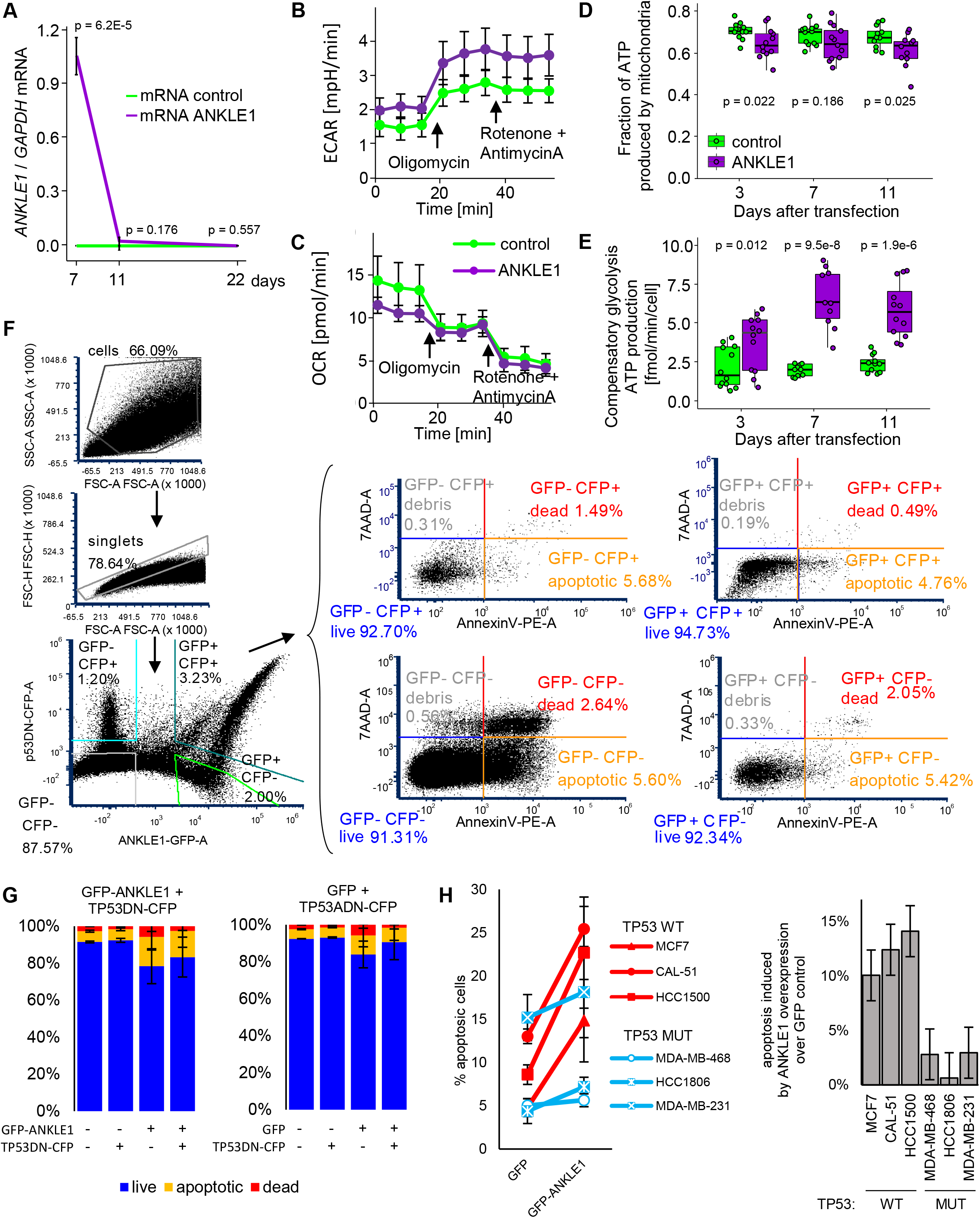
ANKLE1 induces mitophagy, which results in a change in energy production and apoptosis resistance that is dependent upon TP53 inactivation. A) *ANKLE1* mRNA recovers to pre-transfection levels 11 days after ANKLE1 overexpression. B-C) ECAR (B) and OCR (C) measurements obtained during the ATP rate assay in ANKLE1-overexpressing cells show higher extracellular acidification rate and lower oxygen consumption rate compared to control GFP cells. This observation is consistent with switching of the metabolism from oxidative phosphorylation to glycolysis as a main energy source. D) ANKLE1 decreases the fraction of ATP produced by mitochondria in HEK293T cells. E) ANKLE1 increases maximum compensatory glycolysis in HEK293T (p-values are calculated with a two-tailed t-test). F) An example gating strategy isolates GFP-ANKLE1 (or control) and CFP-TP53 to assess the level of apoptosis in cells overexpressing ANKLE1 and/or a TP53 dominant negative mutant. G) The level of apoptosis in HEK293T is not affected by ANKLE1 overexpression, presumably because TP53 is already inactive in HEK293T cells. H) ANKLE1 overexpression specifically increases apoptosis in p53 wild-type cell lines (MCF7, HCC1500 and CAL51) compared to p53-mutant breast cancer cell lines (HCC1806, MDA-MB-231 and MDA-MB-468). These results support our conclusion that ANKLE1 causes apoptosis in presence of wild-type p53, and not in presence of mutant p53.

**Fig. S7.**
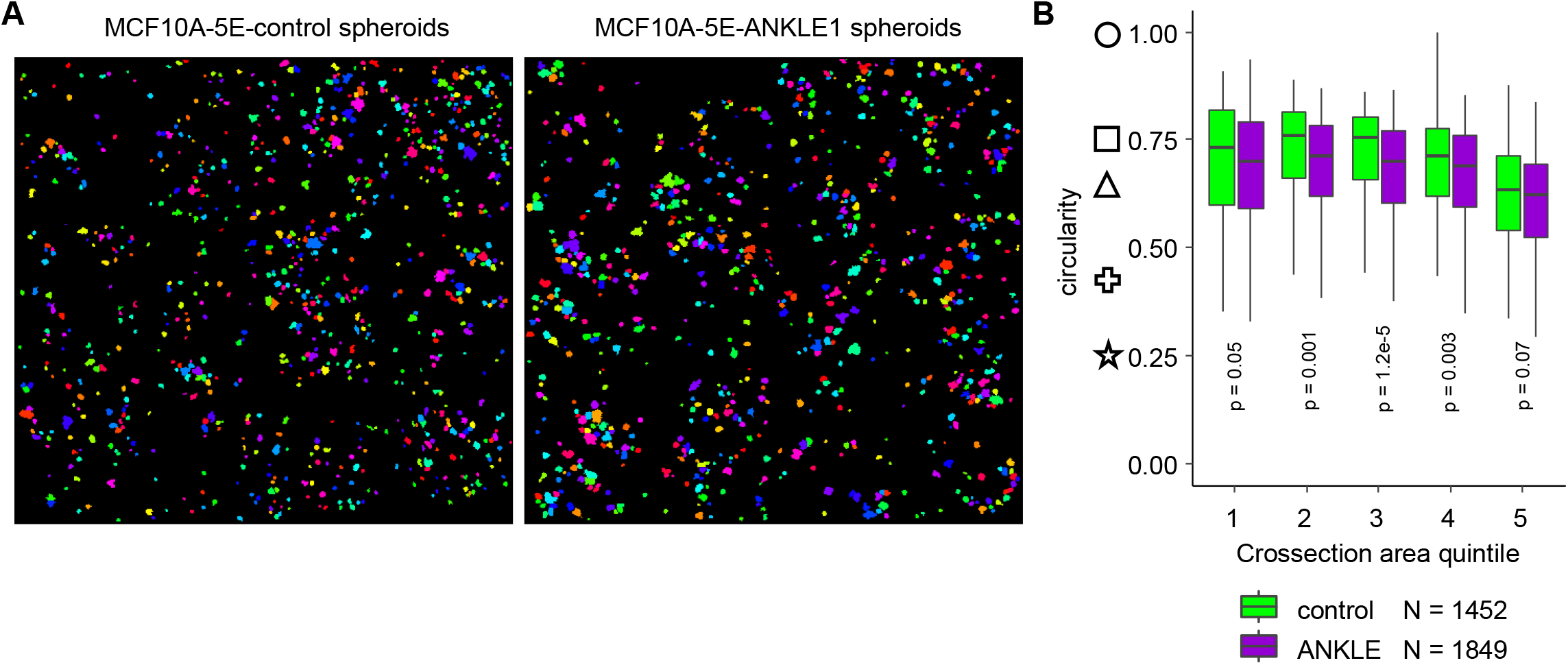
ANKLE1 induces distortion of MCF10A-5E spheroids, irrespective of spheroid size. A) Images that were quantified by OrganoSeg software are distinguishable as individual spheroids. B) We separated spheroids into quintiles based upon their circularity. Note that quintiles are defined by the pooling of control and ANKLE1 cells, so the ranges of adjacent box plots are identical. These results confirm that while larger spheroids tend to be less circular, lower circularity among ANKLE1-expressing spheroids is not driven by their larger size (p-values are calculated with a two-tailed t-test).

